# Leveraging Lineage Barcodes as Natural Augmentations for Contrastive Learning of Cell Fate in scRNA-seq Data

**DOI:** 10.1101/2024.10.28.620670

**Authors:** Shizhao Joshua Yang, Yixin Wang, Kevin Z. Lin

**Author notes:** Correspondence to: Shizhao Joshua Yang < >, Kevin Z Lin < >.

## Abstract

Deciphering how cells commit to future fates is essential for developing precision therapeutics that can reprogram stem cells or modulate immune functions. However, isolating these fate-determining signals in single-cell lineage tracing (scLT) remains challenging because differentiation programs are often confounded by unrelated processes like the cell cycle. To address this, we introduce Lineage-aware Contrastive Learning (LCL), a framework that treats inheritable lineage barcodes as a “natural” data augmentation to isolate subtle, lineage-specific signals. LCL utilizes a semi-supervised architecture to align unlabeled cells, facilitating the transfer of lineage structures to clinical datasets where explicit barcoding is unavailable. We demonstrate LCL’s utility by predicting future cell-type compositions from early-time points, effectively modeling longitudinal fate commitment from cross-sectional data. Benchmarking on hematopoietic and fibroblast systems shows that LCL significantly outperforms standard models like scVI, establishing contrastive learning as a scalable paradigm for understanding and potentially manipulating cellular differentiation.

## 1. Introduction

Understanding how cells commit to their future fates – the transition from an initial state to specific final states – enables therapeutics to precisely manipulate undifferentiated cells. Such knowledge is critical for improving immunotherapies (Sugiura et al., 2022) and reprogramming protocols (Sharma et al., 2022). However, the destructive nature of single-cell sequencing prevents direct longitudinal study of individual cells. To overcome this, single-cell lineage tracing (scLT) protocols infect cells with inheritable lentiviral barcodes; as cells divide, their progeny inherit these markers, allowing scientists to track ancestry relationships over time (Chen et al., 2022a; Jones et al., 2023). These protocols simultaneously sequence gene expression and lineage barcodes, providing richer context than standard single-cell RNA-sequencing (scRNA-seq) (Wagner & Klein, 2020). The resulting combination of temporal and clonal information offers unprecedented insights into cell fate dynamics (Fig. 1). However, existing computational methods struggle to isolate subtle, fate-determining signals from other biological confounding processes.

**Figure 1.**
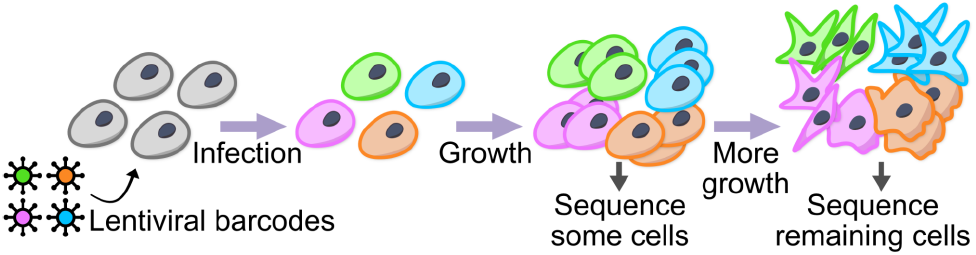
Schematic of single-cell lineage barcoding protocol.

Many methods have been developed to model scRNA-seq data and understand fate commitment without using lineage barcodes (Trapnell et al., 2014; Street et al., 2018; Setty et al., 2019; Bergen et al., 2020; Chen et al., 2022b). However, all these approaches share a fundamental obstacle. A cell’s gene expression is a composite of many mechanisms that occur simultaneously, not all of which are related to the lineage-specific developmental programs of interest. For example, gene expression gives only partial information about the cell cycle, cell identity, and various signaling mechanisms. These factors all confound the subtle signals that indicate differentiation and future fate (Pascual-Ahuir et al., 2020; Eisenstein, 2020; Nitzan & Brenner, 2021). Although lineage barcodes (i.e., labels) reveal critical biological findings that are statistically invisible to standard methods, the vast majority of currently available single-cell datasets lack these explicit markers. To address this, we develop Lineage-aware Contrastive Learning (LCL). LCL is a semi-supervised deep learning method with a contrastive architecture. It treats lineage labels as “natural” data augmentations to isolate fate-determining signals. Crucially, LCL enables transfer of learned lineage labels from scLT datasets to non-barcoded scRNA-seq datasets. By aligning the embedding spaces of both datasets, LCL helps researchers model cellular fates in typical scRNA-seq studies. This is especially useful where in vitro barcoding is unavailable.

## 2. Related work

The challenge in isolating the lineage signals that drive different cell fates is that these signals are often overshadowed by more prominent signals, such as cell-type variation, which may be orthogonal to cell fate in complex biological systems. See Fig. 2 for the schematic of scLT data. Many current methods in single-cell analysis use variational autoencoder (VAE) architectures to model single-cell data (Lopez et al., 2018; Choi et al., 2023; Weinberger et al., 2023; Boyeau et al., 2023), which excel at reconstructing gene expression. However, since these methods are unsupervised, the resulting representations may not sufficiently isolate fate commitment signals from other biological processes in the cell, making it difficult to study how cells commit to their fates.

**Figure 2.**
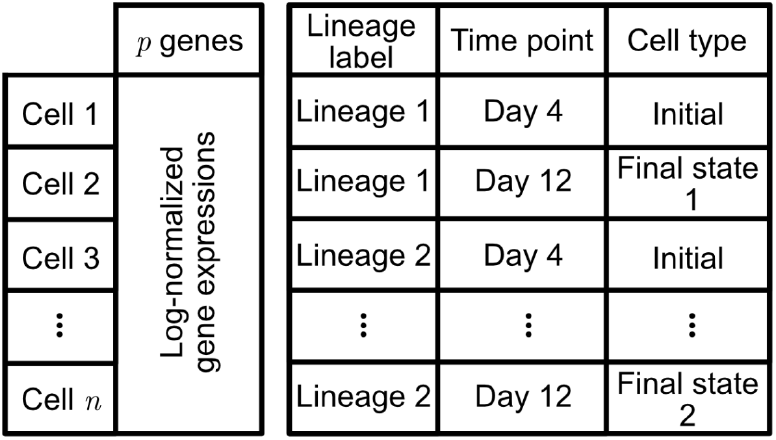
Schematic of a scLT dataset containing gene expression vectors, lineage, time point, and cell type of each cell.

Since we want to model the lineage barcodes, our problem is *supervised*. On the other hand, standard deep learning prediction methods are also not suitable for learning how gene expression encodes fate programs because the signal-to-noise ratio is too low. For instance, using VariancePartition (Hoffman & Schadt, 2016) to quantify how much measured covariates explain the variability of each gene’s expression, we see that while some genes strongly encode a cell’s lineage, the vast majority of genes are dominated by variability in cell type or time point (Fig. 3). Therefore, we deploy contrastive learning to uncover the faint gene expression signals specific to lineage information. Contrastive learning learns an embedding in which cells in the same lineage are close to each other if and only if they are in the same lineage. This idea originates from the computer vision literature to embed images where *data augmentation* mechanisms (such as rotations, cropping, etc.) of individual images, which do not fundamentally change the content of the image, help the deep learning model to learn more generalizable representations (Chen et al., 2020; Chuang et al., 2020; Khosla et al., 2020). By treating lineage barcodes as a naturally occurring data augmentation mechanism, we encourage the deep learning model to hone in on gene expression components specific to lineage information. In what’s to come, we compare our proposed method to CoSpar (Wang et al., 2022) and GEMLI (Eisele et al., 2024), two other methods that specifically incorporate the lineage barcodes into their statistical modeling. See Appendix S2 for a further discussion of related work.

**Figure 3.**
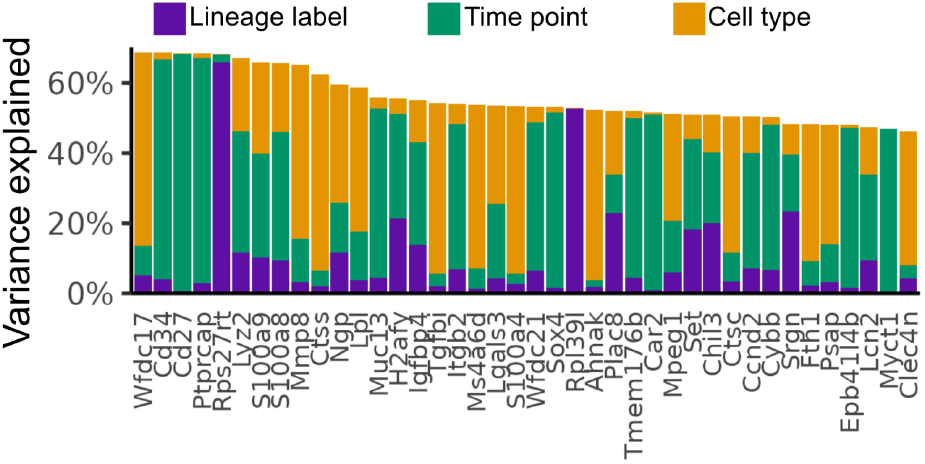
The amount of variance explained by the lineage label, time point, and cell type among the 40 genes with the greatest variance in our hematopoietic differentiation system (Weinreb et al., 2020). Most gene expression variability is explained by cell type or time point, not by lineage labels.

## 3. Method

### 3.1. LCL’s semi-supervised contrastive learning architecture and loss function

The primary objective of Lineage-aware Contrastive Learning (LCL) is to learn a representation space that prioritizes lineage fate-determining signals over dominant transcriptomic noise. Given a dataset 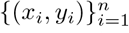 of *n* cells with *p*-dimensional gene expression *x*_*i*_ and lineage labels (i.e., barcodes) *y*_*i*_, we learn an embedding *z*_*i*_ that disentangles fate-relevant biological programs from confounding processes. Crucially, LCL is not a classifier; its goal is not to predict a cell’s lineage label *y*_*i*_, but rather to learn an embedding that reveals the “fate commitment” of cells as they transition from an initial state to specific final states, even in datasets where lineage barcodes were never explicitly recorded. We optimize the embedding according to the following three goals:

- **Goal #1**: Cluster cells by lineage labels to isolate subtle, fate-determining signatures from confounding sources of variation.
- **Goal #2**: Capture generalizable latent features that allow cells from unseen lineages to naturally cluster by ancestry in the embedding space, despite the model having no prior exposure to those specific labels.
- **Goal #3**: Facilitate the mapping of non-labeled datasets into this lineage-aware embedding, enabling the discovery of lineage fate dynamics in studies where *in vitro* lineage barcoding is impossible.

To achieve these goals, LCL comprises four main components: the cell-pair generator, the base encoder, the projection head, and the loss function (Fig. 4). To enable the model to generalize beyond lineage-labeled cells and transfer its learned representations to unlabeled but related datasets, we introduce a complementary unsupervised part alongside the supervised (contrastive) part. This semi-supervised design allows LCL to leverage both labeled cells (i.e., cells with lineage labels) and unlabeled cells (i.e., cells without lineage labels) during training, thereby enhancing robustness and facilitating lineage-label transfer, as we describe below.

**Figure 4.**
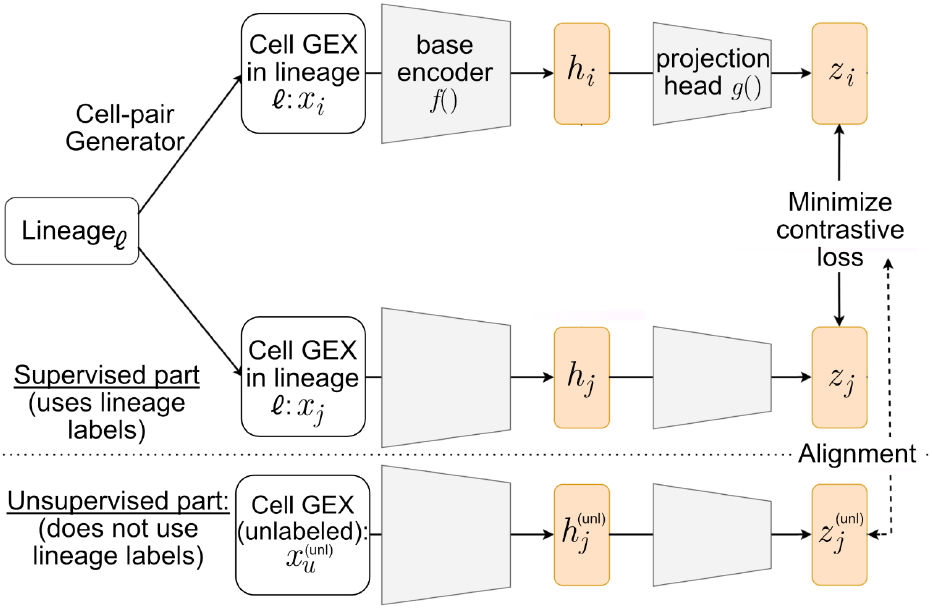
The architecture of LCL uses a contrastive loss on the supervised part, where the lineage labels provide a natural data augmentation mechanism. The cell-pair generator forms positive pairs from the gene expression vectors (GEX) *x*_*i*_ and *x*_*j*_ of two cells with the same lineage label *𝓁*. The unsupervised part leverages unlabeled cells (i.e., no lineage labels), and the alignment penalty ensures the unlabeled cells’ embeddings are well aligned with the labeled cells’ embeddings.

#### Cell-pair generator

For the *supervised part*, most contrastive learning frameworks, such as SimCLR (Chen et al., 2020) and SupCon (Khosla et al., 2020), are designed for images, where data augmentation generates positive pairs of images – two images that are numerically distinct but conceptually the same object. In contrast, our method leverages the inherent properties of lineage-barcoded data to define positive pairs – two cells are considered positive pairs if they belong to the same lineage (i.e., *y*_*i*_ = *y*_*j*_ for two cells *i* and *j*), while cells from different lineages are treated as negative pairs (i.e., *y*_*i*_ ≠ *y*_*j*_). The cell-pair generator first generates all positive pairs used during training. For a specific lineage *L*_*k*_ ∈ {1, …, *L*}, its total number of positive pairs is given by 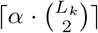, where we call α ∈ (0,1] the size factor. The cell-pair generator then assigns the positive pairs to different training batches by selecting *N* cell pairs from *N* distinct lineages per batch. Within each batch, any two cells from different lineages are deemed as a negative pair. We then design LCL’s remaining components to maximize the agreement between cells from positive pairs (i.e., cells from the same lineage) and minimize the agreement between negative pairs (i.e., cells from different lineages).

For the *unsupervised part*, we extend this framework to include *m* unlabeled cells, 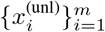. In each training iteration, *U* cells are randomly sampled from the unlabeled dataset. As we will describe, we leverage these unlabeled cells to allow the model to generalize beyond the lineage-barcoded population.

#### Base encoder

The base encoder *f* (·) is a neural network that derives an intermediary embedding *h*_*i*_ using *x*_*i*_, the gene expression vector of each cell *i*. This is applied to each cell in the pairs we constructed. By default, we use a 3-layer Multilayer Perceptron (MLP) as the base encoder using the Rectified Linear Unit (ReLU) activation function, and batch normalization is applied after each layer to normalize the output. The input and output dimensions of *f* (·) are *p* and 64, respectively.

#### Projection head

The projection head *g*(·) consists of a two-layer MLP with ReLU that transforms *h*_*n*_ into a lower-dimensional embedding, denoted by *z*_*n*_ = *g*(*f* (*x*_*n*_)), where the contrastive loss is applied. In standard contrastive learning frameworks, the projection head is often discarded after training and downstream analyses are performed using the base encoder representation (Gupta et al., 2022). In LCL, both the base-encoder embedding *h*_*n*_ = *f* (*x*_*n*_) and the projection-head embedding *z*_*n*_ = *g*(*f* (*x*_*n*_)) can be useful for downstream analyses. The projection-head embedding is directly optimized by the contrastive objective and provides a compact lineage-aware representation, while the base-encoder embedding may retain additional biological variation that can benefit some downstream prediction tasks. Therefore, our open-source package outputs both embeddings, and we report both variants in our empirical evaluations when relevant. The output dimension of *g*(·) is 32.

#### Loss function

For each batch containing *N* labeled cell pairs (i.e., 2*N* labeled cells), we define the loss for a positive pair (*i, j*) using the normalized temperature-scaled cross-entropy loss, a contrastive objective introduced in prior work (Chen et al., 2020),

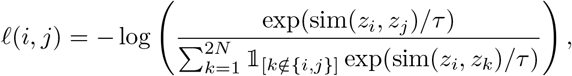

where sim(*z*_*i*_, *z*_*j*_) denotes the cosine similarity ((*z*_*i*_)^*T*^*z*_*j*_)*/*(∥*z*_*i*_ ∥_2_ *z*_*j*_ ∥_2_) and *τ >* 0 is the temperature parameter. The supervised contrastive loss for the labeled cells to achieve Goal #1 is then computed as the average across all positive pairs,

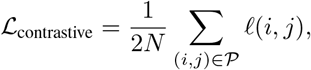

where 𝒫 is the set of positive (same-lineage) cell pairs. This loss term encourages cells of the same lineage to be close together in the embedding space.

#### Semi-supervised penalty for unlabeled cells

To learn robust lineage-informative embeddings (Goal #2) and align the embeddings of unlabeled cells with the lineage-informed space learned from labeled cells (Goal #3), we introduce an additional entropy-based penalty term. For each training iteration, *U* unlabeled cells pass through the same base encoder *f* (·) and projection head *g*(·) to obtain embedding vectors 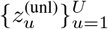. We design the following penalty term to encourage alignment between the labeled and unlabeled embedding spaces. Importantly, this penalty should be interpreted as manifold alignment rather than strict lineage classification. Although unlabeled cells are softly compared to labeled cells in each mini-batch, they are not assigned fixed discrete lineage labels. Instead, the penalty encourages unlabeled cells to align with confident regions of the lineage-informed transcriptomic manifold. Thus, in our biological setting, generalization means mapping unseen lineages onto shared functional cell-state structure, rather than creating a separate orthogonal cluster for every new barcode. Specifically, each unlabeled cell forms a soft assignment to the 2*N* labeled projections using temperature-scaled cosine similarity,

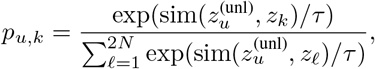

for *k* ∈ {1, &, *N*}. We then apply an entropy penalty that encourages confident assignments (i.e., low-entropy) between unlabeled cells and nearby labeled neighbors,

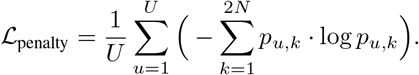

The entropy penalty reduces the embedding gap between labeled and unlabeled cells.

The full training objective combines the supervised contrastive loss and the unsupervised entropy penalty:

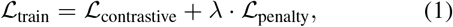

where *λ >* 0 controls the strength of the semi-supervised regularization. This joint minimization of ℒ_train_ improves the model’s robustness on unseen data and allows cells without labels to be mapped to probable lineage neighborhoods in the learned space. As we empirically show, this regularization yields smoother cluster boundaries and lower KNN test error across both labeled and unlabeled datasets.

### 3.2. Mini-batch construction and semi-supervised sampling

LCL is trained using lineage-based contrastive learning, optionally augmented with a semi-supervised penalty. Training is organized around a cell-pair generator, denoted 𝒞(*α*), which constructs lineage-based positive cell pairs and assembles them into mini-batches (see Appendix S3 for details).

For each mini-batch, 𝒞(*α*) samples *N* positive cell pairs from distinct lineages whenever possible, yielding 2*N labeled cells*. These labeled cells are used to compute the supervised contrastive loss, where positive and negative relationships are defined entirely by lineage membership. In addition, when semi-supervision is enabled, 𝒞(*α*) independently samples *U unlabeled cells* that do not participate in positive or negative pair construction. Instead, these unlabeled cells contribute only through an entropy-based penalty, which encourages their embeddings to align with the lineage-informed structure learned from the labeled pairs. Unlabeled cells may be drawn either from an external unlabeled dataset or from a held-out unlabeled pool within the same dataset when no external unlabeled data are available. In both cases, unlabeled cells are never used to define supervised contrastive labels and do not affect the construction of positive pairs.

Details on data preprocessing, training, scalability, and diagnostics are provided in Appendix S3 and Appendix S5, where we further document the role of hyperparameters such as the number of positive pairs *N*, the number of unlabeled cells *U* per mini-batch, the temperature *τ*, the size factor *α*, and the semi-supervised regularization weight *λ*.

### 3.3. Downstream application: Identify genes with lineage information via integrated gradients

Lineage barcodes serve as labels that enable LCL to separate heritable, fate-relevant transcriptional programs from dominant but potentially distracting sources of variation (e.g., cell type or time effects). After training, we aim to make the learned lineage-aware embedding interpretable by identifying which genes most strongly drive a cell to reside in its lineage-specific region of the embedding space. Concretely, we treat genes as “lineage-informative” if changing their expression most increases a cell’s alignment with its lineage cluster in the LCL embedding, yielding a ranked gene list that can be inspected for biological plausibility and used for downstream hypothesis generation.

To score gene contributions, we apply Integrated Gradients (IG) to a scalar function that measures how well a cell’s em-bedding aligns with its lineage centroid (Sundararajan et al., 2017). For each lineage *𝓁*, we compute its centroid *c*_*𝓁*_ as the mean of the LCL embeddings *z*_*i*_ of cells in that lineage, where *z*(*x*) = *g*(*f* (*x*)) is the (normalized) projection output used throughout our downstream analyses. We then define *F*_*𝓁*_ (*x*) = cos(*z*(*x*), *c*_*𝓁*_) and compute IG attributions for each gene *j* relative to a baseline *x*^*′*^ (zero or mean expression): 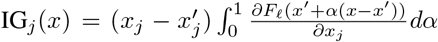 approximated numerically. We summarize lineage-level gene importance by averaging |IG_*j*_| across cells within a lineage, and obtain a global ranking by averaging these lineage-level scores across lineages. See Appendix S4 for more details.

### 3.4. Downstream application: Predict the future composition of lineages

To quantify cellular fate commitment, we use the LCL embedding of early time point cells to predict the future cell-type composition of their respective lineages. For a system with *c* terminal states, we represent the fate of lineage *i* as a *c*-dimensional proportion vector, where each entry denotes the relative frequency of a specific cell type within that lineage at the final time point. We train a linear decoder that maps individual cell embeddings to logits over terminal cell types, followed by a softmax to obtain a predicted composition distribution.

While standard cross-entropy is well suited for predicting discrete, hard-target classes of individual cells, our goal here is to predict the future composition of an entire lineage. Because each lineage’s fate is represented as a soft target probability distribution over multiple terminal cell types, we optimize and evaluate the linear decoder using KL divergence, which naturally measures the discrepancy between predicted and observed compositional distributions.

This approach allows us to evaluate whether the learned embeddings capture latent, fate-determining signals that drive longitudinal differentiation from cross-sectional data.

## 4. Experimental Setup

### 4.1. Datasets

We briefly introduce the datasets in our study; more details are available in Appendix S4. In all our analyses, we first perform feature selection to analyze only the top *p* = 2000 highly variable genes, and we log-normalize the gene expression vectors 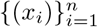, as is standard in single-cell literature.

- **Hematopoietic Differentiation** (Weinreb et al., 2020): The authors designed the LARRY protocol to barcode hematopoietic stem cells in mice over a 6-day experiment to study how cells commit to differentiating into certain cell types. We will call this the “LARRY dataset,” which contains 41,093 cells across 11 cell types, with 2,813 lineages, sampled at 3 different time points.
- **Fibroblast Reprogramming** (Biddy et al., 2018): The authors designed the CellTagging protocol to barcode mouse embryonic fibroblasts when reprogramming them into induced endoderm progenitors (iEPs). Cells in certain lineages failed to reprogram (i.e., “dead-end”). We will call this the “CellTag dataset,” which contains 6,534 cells with 169 different lineages, across 3 main cell types sequenced at various time points in a 28-day experiment (6 time points).
- **Followup Fibroblast Reprogramming** (Jindal et al., 2024): The authors improved the CellTagging protocol to the CellTag-multi protocol and uses this to further study the fibroblast-to-iEP reprogramming system. We will call this dataset the “CellTag-multi dataset,” which contains 22,238 cells across 1,367 lineages, spanning 5 main cell types in a 21-day experiment (3 time points).

### 4.2. Simulation via pseudo-real data

To evaluate LCL across varying signal-to-noise ratios, we simulate lineages using the LARRY dataset by reassigning labels based on a subset of gene expression values. We define a difficulty parameter *β* ∈ {0.1, 0.3, &, 0.9}, representing the top fraction of highly variable genes used to induce lineage structure. After ranking genes by variance, we project the top *β*-fraction onto its principal components and define simulated lineages via Leiden clustering. Lower *β* values yield benchmarks that are more challenging, with sparse lineage information, ensuring that methods like scVI cannot trivially recover the structure via standard transcriptomic clustering. Five trials are conducted for each *β*; see Appendix S4 for more details.

### 4.3. Evaluation metrics and baseline methods

After fitting LCL, we evaluate the quality of the learned lineage-aware embedding using various qualitative and quantitative methods.

#### Evaluation for training cells

To evaluate global lineage coherence, we use the Calinski-Harabasz (CH) Index (Cali ński & Harabasz, 1974) to measure cluster separation and cohesion. Additionally, we employ Gene Expression Memory-based Lineage Inference (GEMLI) (Eisele et al., 2024) to assess the predictive power of the latent space. GEMLI trains a random forest classifier on latent cell-pair representations to predict whether two cells originate from the same lineage; we report performance using the Area Under the Precision-Recall Curve (AUPRC) and the Area Under the Receiver Operating Characteristic curve (AUROC). See Appendix S3 for details.

#### Evaluation on held-out test cells

We evaluate how well LCL’s embedding generalizes beyond the cells used for train-ing, which is critical for assessing robustness to overfitting and the extent to which the embedding captures biologically meaningful structure. To this end, we split cells into disjoint training and test sets prior to model fitting. All evaluation metrics are computed exclusively on the held-out test cells, which are not used to train either the LCL encoder or any downstream predictors.

We consider two complementary evaluation metrics. First, we assess whether cells from the same lineage remain close in the embedding space by training a *K*-nearest neighbor (KNN) classifier on the training-set embeddings and corresponding lineage labels, and evaluating its classification accuracy on the test-set embeddings. Throughout this work, we use *K* = 5. High test accuracy indicates that lineage structure learned during training generalizes to unseen cells.

Second, we evaluate whether the learned embedding preserves information predictive of future cell fate, as described in Section 3.4. Using only the training split, we fit a linear softmax decoder that maps early time-point cell embeddings to lineage-level future cell-type composition vectors. Decoder training and early stopping are performed exclusively on the training split. Performance is then evaluated on the held-out test cells by computing the Kullback–Leibler (KL) divergence between the predicted and observed composition distributions. Lower KL divergence on the test set indicates that LCL successfully isolates latent fate-determining signals that generalize beyond the training data.

#### Benchmarking baselines and visualization

We compare LCL against scVI (Lopez et al., 2018), Supervised UMAP (McInnes et al., 2018), cross-entropy, and triplet-loss embedding baselines. For qualitative assessment, we visualize embeddings using UMAP; for quantitative assessment, we use the KNN lineage-prediction and KL-divergence composition-prediction metrics described above. Implementation details for all baseline methods are provided in Appendix S4.8.

## 5. Results

### 5.1. LCL reliably extracts the lineage information on pseudo-real data

In our first set of results, we leverage our simulated scLT datasets to confirm LCL’s ability to extract lineage information at varying signal strengths. Using the pseudo-real datasets with varying *β*, we see that LCL consistently performs well across all difficulty levels (Fig. 5). The high CH Index shows this across all five trials’ *β* values. This demonstrates LCL’s robustness in extracting lineage information, even when the signal becomes harder to distinguish. In contrast, scVI and Supervised UMAP perform worse at extracting the lineage-dependent signal from the gene expression count matrix at all levels of *β*.

**Figure 5.**
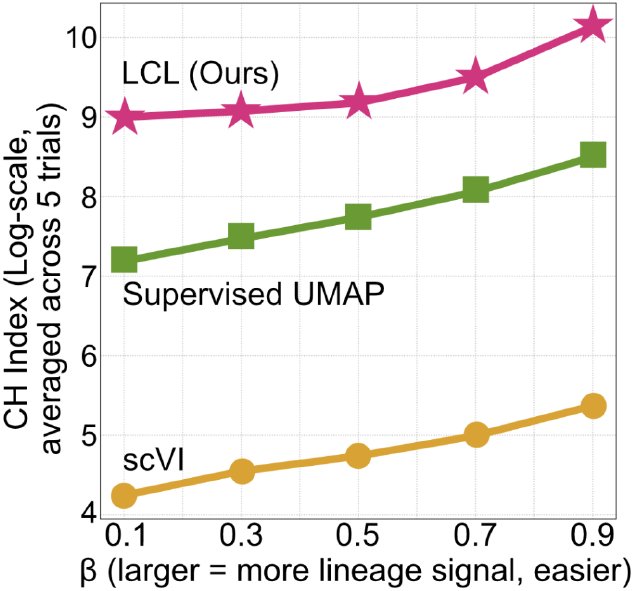
CH Index of different *β*’s comparing how LCL compares with scVI and Supervised UMAP, where a higher index means the embedding better separates the different lineages.

### 5.2. LCL learns lineage-informed embeddings in various biological contexts

After demonstrating LCL’s capabilities on simulated data, we next analyze the LARRY and CellTag datasets for our second analysis. We demonstrate that LCL yields higher-quality embeddings than scVI and Supervised UMAP on both datasets, indicating that cells from the same lineage are more clustered together. Visually, the LCL embeddings form denser clusters in the UMAP, where we observe that the cells from the same lineage form “islands” (Fig. 6B). In contrast, Supervised UMAP shows a visual trend towards lineage clustering but still exhibits significant overlap between cells from different lineages (see Appendix Fig. S8). Meanwhile, scVI embeddings demonstrate a mixture of cells from various lineages without clear separation, likely due to its unsupervised nature (Fig. 6A). These findings are also supported qualitatively by the CH Index, which shows that LCL is more effective at separating cells by lineage and, therefore, better isolates lineage-related information. See Appendix S5 for more plots and results.

**Figure 6.**
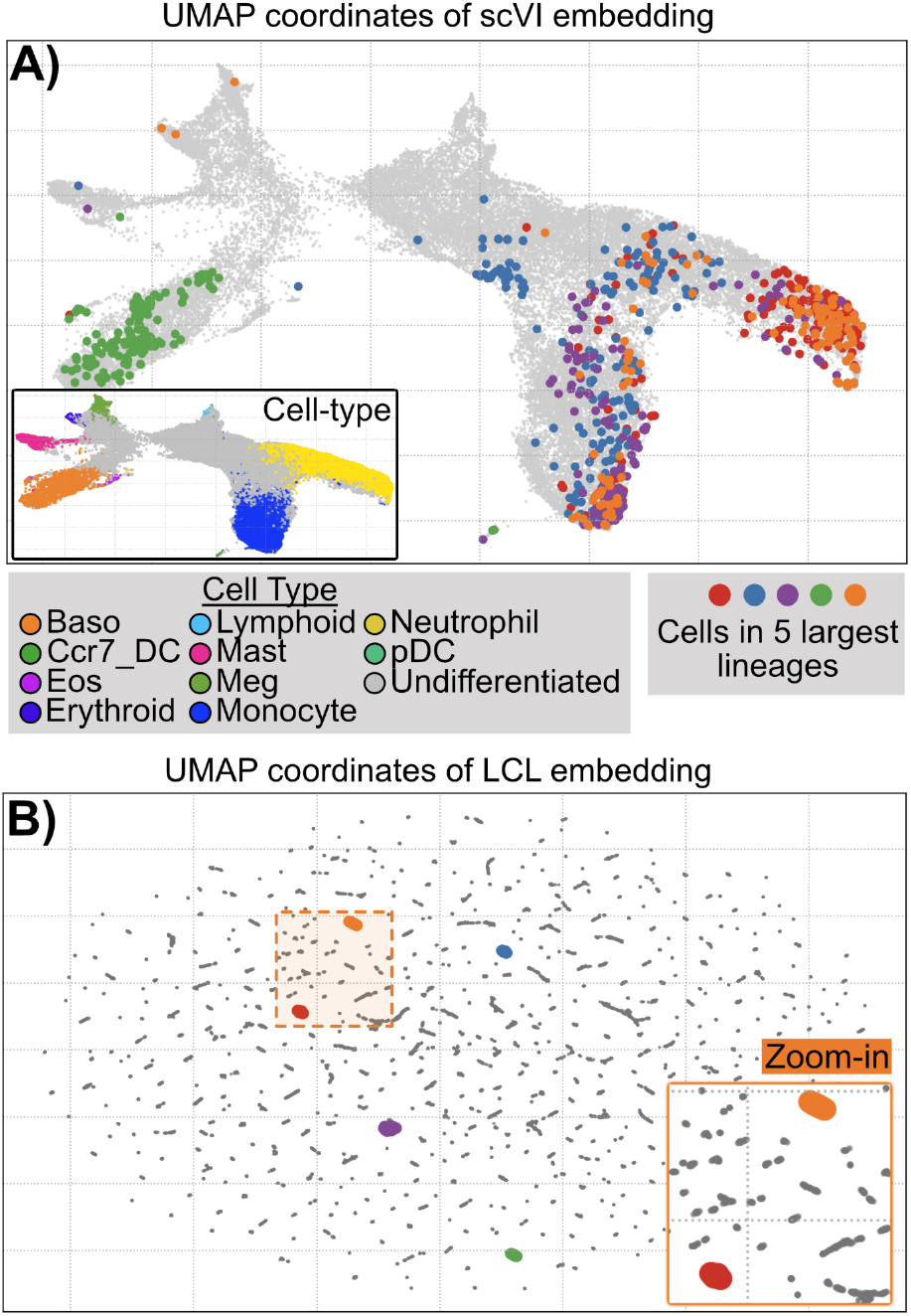
**A)** UMAP of the scVI embedding for the LARRY dataset. The inset shows the cells colored by cell type. **B)** UMAP of the LARRY embeddings. The inset shows a zoom-in of the orange lineage. In both UMAPs, the cells in the five largest lineages are highlighted in different colors in their respective datasets. Surprisingly, although the LCL embedding visually shows “islands,” the embedding performs well on the test set.

One might question whether the island-like formations in LCL’s UMAP indicate overfitting, so we use the KNN classifier to evaluate the generalizability of LCL’s embeddings.

We train a KNN classifier on the training-cell embeddings to predict their lineages and evaluate the test error on held-out test-cell embeddings.

For the CellTag dataset and the largest 200 lineages from the LARRY dataset, LCL embeddings perform competitively across both held-out lineage prediction and future-composition prediction. In particular, the base-encoder embedding gives the strongest overall KNN and KL-divergence performance, while the projection-head embedding also improves future-composition prediction relative to the baseline embeddings (Table 1).

**Table 1.** Performance comparison of embedding methods on lineage prediction and future cell type composition prediction. Lower values indicate better performance. The best values are bolded.

| Dataset | Method | KNN Test Error ↓ | KL Divergence ↓ |
| --- | --- | --- | --- |
| CellTag | scVI | 0.529 | 0.627 |
|  | Supervised UMAP | 0.628 | 0.595 |
|  | CE | 0.437 | 0.771 |
|  | Triplet | 0.530 | 0.606 |
|  | LCL (proj) | 0.459 | 0.414 |
|  | LCL (base) | **0.434** | **0.292** |
| LARRY (Largest 200) | scVI | 0.892 | 0.549 |
|  | Supervised UMAP | 0.941 | 0.903 |
|  | CE | 0.898 | 0.796 |
|  | Triplet | 0.905 | 0.688 |
|  | LCL (proj) | 0.864 | 0.361 |
|  | LCL (base) | **0.858** | **0.335** |

This further confirms the generalizability of LCL’s embed-dings. More comprehensive results, including comparisons with scVI and Supervised UMAP, other definitions of test cells, and the CH Index, are presented in Appendix S5, where LCL consistently achieves higher test accuracy across both datasets. All these results point to somewhat surprising conclusions: although the LCL embedding visually shows “islands,” it performs well on test sets. This suggests that LCL uncovers meaningful biological signals specific to cells within the same lineage, despite their different cell types.

### 5.3. LCL’s embedding enables improved predictions of future cell type compositions

We next ask whether lineage-informed embeddings improve prediction of future cell fate. For each early-time-point cell, we use its embedding to predict the future cell-type composition of its lineage, as described in Section 3.4. Because each lineage’s fate is represented as a probability distribution over terminal cell types, we evaluate prediction accuracy using KL divergence between the predicted and observed future compositions.

Across both the CellTag dataset and the largest 200 lineages from the LARRY dataset, LCL embeddings achieve lower KL divergence than scVI, Supervised UMAP, cross-entropy, and triplet-loss baselines (Table 1). The base-encoder embedding performs best in these experiments, suggesting that it retains additional biological variation useful for composition prediction. These findings highlight LCL’s ability to isolate lineage-specific signals that drive cell-fate decisions, enabling it to outperform other methods in predicting future cell-type compositions.

### 5.4. LCL enables new biological investigations of unlabeled scRNA-seq studies by leveraging lineage-barcoded data

In our final data analysis, we demonstrate that LCL enables the transfer of lineage-informed structure from a lineage-barcoded reference dataset to a related but unlabeled dataset. This setting reflects a common practical scenario in which lineage barcoding is available only for a subset of experiments, while additional datasets measure similar biological systems without explicit lineage annotations. To facilitate this experiment, we use the CellTag and CellTag-multi as the unlabeled and labeled datasets, respectively. Because both datasets capture related biological processes, the learned embedding provides a shared representation space in which lineage-related structure can be transferred. Furthermore, although we treat CellTag as an unlabeled dataset for our experiment, we can use its lineage information to validate LCL’s accuracy in label transfer. See Appendix S4 for the details of how we perform this analysis.

After fitting LCL on the CellTag-multi (labeled) dataset, we assess how the cells in the CellTag (unlabeled) dataset co-localize based on their withheld lineage labels. After integrating the datasets together, we assess suitable integration between the two datasets (Fig. 7A). We don’t expect perfect integration, since the lineages in each dataset are distinct. Additionally, when focusing on the CellTag cells, we observe strong lineage co-localization despite LCL not having used their lineage labels during training (Fig. 7B). This demonstrates LCL’s impressive ability to learn the fate-determining biology of lineages even when no lineages are observed.

**Figure 7.**
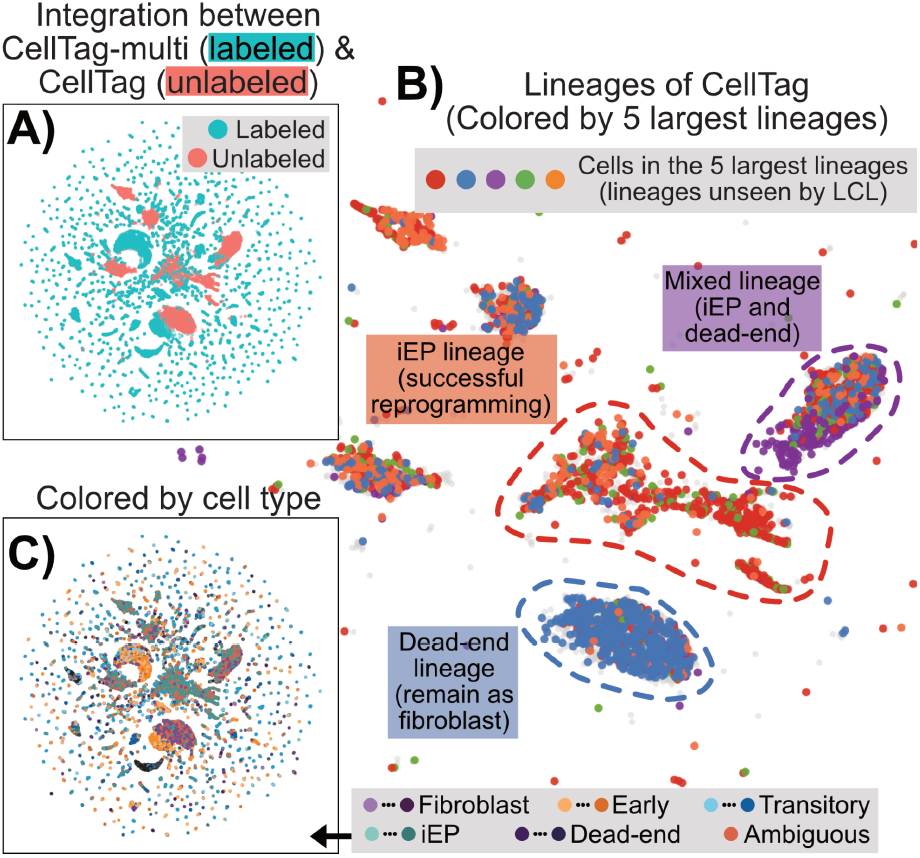
**A)** LCL embedding’s UMAP showing cells from both datasets to demonstrate the integration. **B)** Subset of only the cells from the CellTag dataset (unlabeled), colored by the largest 5 lineages. The shown lineages are not used by LCL and are distinct from the lineages in the CellTag-multi dataset (labeled) used during training. Three of the unseen lineages that tightly clustered are highlighted (“islands”), labeled by predominate fate. **C)** Cells from both datasets colored by cell type.

To evaluate how effectively different embeddings capture lineage-specific signals, we use GEMLI to classify whether two cells are from the same lineage or not. As shown in Table 2, LCL significantly outperforms other methods. Notably, LCL nearly triples the AUPRC of deep generative models (scVI), the log-normalized gene expressions, and a random-guess baseline. These results demonstrate that LCL successfully isolates salient lineage variations from transcriptomic noise, providing a latent representation that is fundamentally more reflective of true fate commitment than standard representation learning approaches.

**Table 2.** AUPRC and AUROC of GEMLI classification on different embeddings. The best values are bolded.

| Metric | LCL | scVI | Log_norm | Random guess |
| --- | --- | --- | --- | --- |
| AUPRC | **0.600** | 0.177 | 0.221 | 0.027 |
| AUROC | **0.825** | 0.601 | 0.637 | 0.5 |

LCL uniquely uncovers critical biological programs that are statistically invisible to alternative methods (Table 3). We compare against CoSpar (Wang et al., 2022), a framework using sparse optimization to infer state transition maps, and VariancePartition (Hoffman & Schadt, 2016). All three methods model the relation between lineage labels and gene expression. While CoSpar and VariancePartition fail to reach significance, LCL successfully identifies the negative regulation of cell differentiation (adjusted p-value = 0.03), capturing the *Wnt4* and *Sfrp1* axis established in prior work as the primary differentiator between successful iEP conversion and dead-end trajectories. Furthermore, LCL resolves TGF-*β* signaling (adjusted p-value = 0.04), endothelial cell migration (adjusted p-value = 0.04, featuring *Bmp4*), and extracellular matrix assembly (adjusted p-value = 0.03, featuring *Col1a2*), all of which are hallmark mesenchymal-to-epithelial transition programs described by the original authors (Biddy et al., 2018; Jindal et al., 2024). These results demonstrate LCL’s superior ability to disentangle subtle, fate-determining signals from high-dimensional scRNA-seq data where existing lineage-aware methods fail.

**Table 3.** Adjusted p-values for pathways of interest across methods. The best values are bolded.

| Pathway | LCL | CoSpar | VP |
| --- | --- | --- | --- |
| Negative regulation of cell differentiation | **0.03** | 0.22 | 0.91 |
| Regulation of endothelial cell migration | **0.04** | 0.82 | 0.74 |
| Regulation of transforming growth factor beta receptor signaling pathway | **0.04** | 0.89 | 0.81 |
| Extracellular matrix assembly | **0.03** | 0.50 | 0.85 |

**Table 4.** Differential expression of LCL clusters. Adjusted p-values, log2 fold-changes, and expressed fate labels are shown.

| Gene | Adjusted p-value | log2FC | Expressed fate |
| --- | --- | --- | --- |
| *Ptn* | 0.00e+00 | 3.25 | Dead-end |
| *Sfrp1* | 0.00e+00 | 3.07 | Dead-end |
| *Shh* | 1.40e-83 | 3.08 | iEP |

Lastly, we explore the biological insights LCL uncovers from the CellTag (unlabeled) dataset, despite not being trained on its lineage labels. This is to reinforce the notion that LCL can help advance the biology of fate commitment of *unlabeled* scRNA-seq data, which is much more abundant than lineage-barcoded scRNA-seq data. We perform high-resolution Leiden clustering of the CellTag cells using the LCL embedding to isolate the different “islands.” Then, we identify marker genes distinguishing these clusters using the Wilcoxon test in a one-versus-rest differential expression analysis. Key genes of the failed reprogramming (i.e., *Ptn, Sfrp1*) and successful reprogramming into iEP fates (i.e., *Shh*) have both significant p-values and high log fold-changes. We are able to validate the fate-specificity of these genes, since CellTag actually has lineage barcodes. We showcase additional results in Appendix S5.

## 6. Discussion

We introduce a novel deep learning framework, Lineage-aware Contrastive Learning (LCL), to address challenges in analyzing scLT data. LCL isolates lineage-specific gene expression signals, which are often overshadowed by other biological processes in conventional methods. By leveraging contrastive learning, LCL effectively disentangles lineage-related signals from cell-type- and time-specific signals, enabling us to predict cell fate with higher accuracy and identify critical fate-determining genes.

We demonstrate that contrastive learning is a promising framework for advancing our understanding of fate commitment in lineage-traced cells. Our results show that LCL outperforms existing methods in clustering lineage-specific signals and predicting future cell-type composition. A limitation of our approach is its reliance on sufficient cell counts per lineage to form robust contrastive pairs. While LCL maintains a moderate correlation (Pearson *r* = 0.3587) between lineage size and prediction accuracy, extremely small lineages naturally exhibit higher variance and lower predictive performance, representing a primary failure mode when applying contrastive augmentations to sparse clonal data. This line of future research can have broad implications for improving our understanding of cellular differentiation and development.

## Code availability

Our LCL architecture and analysis code are available on GitHub (SZ-yang/Lineage_aware_ContraLearn).

Additional details about the code and datasets are in the Appendix S1.

## Acknowledgments

We thank Brandon Hadland, the anonymous ICML reviewers, and other members of the Lin Lab for support, helpful discussions, and feedback on this work. Research reported in this manuscript was supported by the National Institute of General Medical Sciences (NIGMS) of the National Institutes of Health (NIH) under award number: R35GM162089.

## Impact statement

This paper introduces a machine learning framework designed to improve the interpretation of single-cell lineage tracing data. By isolating subtle, fate-determining signals from complex transcriptomic noise, our work advances the biological understanding of cellular differentiation and development. These advancements have significant potential societal benefits in the field of health and medicine, specifically by facilitating the design of precision therapeutics, improving immunotherapy outcomes, and refining stem cell reprogramming protocols.

