## Appendix for "Leveraging Lineage Barcodes as Natural Augmentations for Contrastive Learning of Cell Fate in scRNA-seq Data"

### A. Data Availability

The LARRY data (Weinreb et al., 2020) is publicly available and can be downloaded at https://github.com/AllonKleinLab/paper-data/blob/master/Lineage_tracing_on_transcriptional_landscapes_links_state_to_fate_during_differentiation/README.md. The original count data can be additionally downloaded from https://figshare.com/ndownloader/files/37028569, (which is linked from https://github.com/pinellolab/pyrovelocity/ (Qin et al., 2022)). The CellTag data (Biddy et al., 2018) is publicly available and can be downloaded at https://cellrank.readthedocs.io/en/latest/api/_autosummary/datasets/cellrank.datasets.reprogramming_morris.html#cellrank.datasets.reprogramming_morris. The CellTag-multi data (Jindal et al., 2024) is publicly available and can be downloaded at https://figshare.com/projects/CellTag-multi_2023_DataShare/173508, with additional context and files provided with the help of the Morris lab. The melanoma sequential-treatment dataset from Schaff et al. (Schaff et al., 2024) is publicly available through GEO under accession GSE253739 and through the accompanying Figshare project at https://figshare.com/projects/Clonal_differences_underlie_variable_responses_to_sequential_and_prolonged_treatment/192653.

### B. Further discussion of contrastive learning literature in relation to LCL

We make some remarks to clarify the scope of our paper relative to the literature. First, our notion of "contrastive" differs from Contrastive Language-Image Pre-Training (CLIP) (Radford et al., 2021) for unsupervised learning, or from methods dedicated to finding biological differences between foreground and background datasets, such as contrastiveVI (Weinberger et al., 2023). Second, we focus on static lineage barcodes here (Weinreb et al., 2020; Biddy et al., 2018), as opposed to mutating lineage barcodes, which capture cell phylogenies that are more complex to model (Spanjaard et al., 2018; Simeonov et al., 2021). Lastly, we note that many methods that learn developmental trajectories from scRNA-seq often use scLT data to validate their method and findings (Qin et al., 2022; Mages et al., 2023; Sha et al., 2024). These existing methods primarily aim to separate cells along the leading axes of variation, often leading to separation by cell type (see Figure 3 in the main text). In contrast, our goal in this paper is to specifically model the lineage barcodes to learn an embedding of single-cell data that enables downstream fate-commitment analyses (see Figure 1).

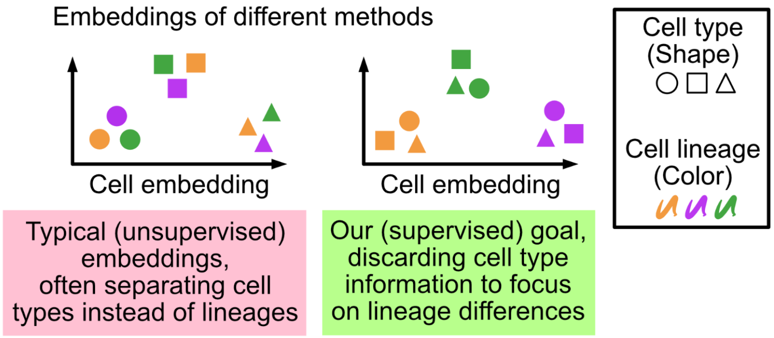


*Figure 1.* Schematic illustrating the differences between conceptual goal in our paper (right, green) compared to most other papers that model scRNA-seq data (i.e., not leveraging lineage barcodes; left, red).

We note that while contrastive learning has been used to embed scRNA-seq data before (Ciortan & Defrance, 2021; Han et al., 2022; Yang et al., 2022), we are, to our knowledge, the first to explore this framework on scLT data, treating the lineage barcodes as data augmentations.

### C. Details of LCL

In Section C.1, we describe more details about the training, the formation of the test set, and the hyperparameters that dictate LCL's performance. In Section C.2, we document how the cell-pair generator (i.e., constructing positive pairs from lineage labels and including unlabeled cells) works. In Section C.3, we provide the pseudocode for LCL. In Section C.4, we document how the training and testing cells are partitioned, when there are both labeled and unlabeled cells involved during training.

#### C.1. Details about training, architecture, and hyperparameters.

**Training.** The dataset is split into training, validation, and test sets. The validation set is used for early stopping to prevent overfitting – if the normalized temperature-scaled cross-entropy loss (NT-Xent) loss on the validation set stops improving after a set number of epochs, training is halted. We optimize the base encoder and projection head neural networks using the AdamW optimizer.

**Forming Test Set.** We create two types of test sets to evaluate different aspects of LCL's generalization ability. The first test set includes 10% of the cells from each lineage, meaning it contains lineages seen during training. The second, more challenging set withholds 10% of the entire lineages, testing the model's ability to generalize to unseen lineages and its potential applicability to datasets that do not have lineage barcodes.

For test sets constructed in the former method, this set is used in the evaluations described in the "Evaluation when unlabeled cells with shared lineages with the labeled cells" section. Specifically, as illustrated in Figure 2, while the training cells are used to fit LCL's embedding and downstream classifiers, the test cells are used to evaluate the accuracy of the classifiers.

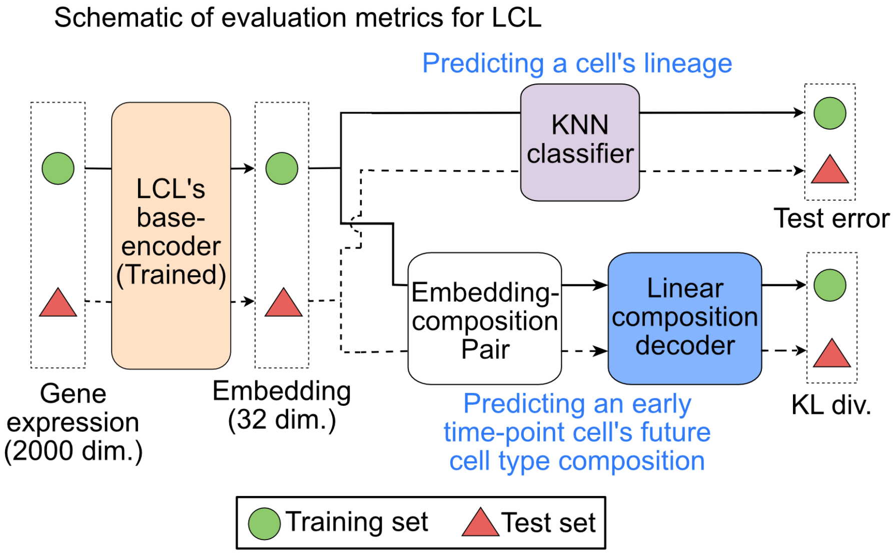


*Figure 2.* Schematic illustrating two evaluation metrics we apply to the LCL embedding. One is for predicting a cell's lineage (top), and another is for predicting an early time-point cell's future cell type composition (bottom).

**Choosing Hyperparameters.** The LCL model has three primary hyperparameters: the size factor $\alpha$, the batch size $N$, and the temperature $\tau$. Size factor $\alpha \in (0, 1]$ and batch size $N$ are both used in the cell-pair generator step. A larger $\alpha$ would yield more pairwise comparisons within and between lineages during training, leading to a better-generalizing embedding at the expense of a higher computational burden. A larger batch size $N \in \mathbb{Z}_+$ means more negative examples are in each batch, facilitating faster convergence and improving performance, especially with fewer training epochs. We use $\alpha = 0.3$ and $N = 100$ as default, but vary them depending on computing resources. The temperature $\tau > 0$ in the NT-Xent loss function controls how the model weights the negative pairs. A small temperature sharpens similarity scores, focusing on harder negatives, which can enhance learning but risks instability or overfitting. In contrast, a larger temperature smoothens the distribution, reducing the emphasis on harder negatives, which may slow learning or yield less distinct embeddings. We find that $\tau = 0.5$ is an appropriate choice for analyzing scLT data.

#### C.2. Details on cell-pair generator $\mathcal{C}(\cdot)$.

We denote the cell-pair generator as $\mathcal{C}(\alpha)$, where $\alpha$ is the size factor. The role of $\mathcal{C}(\alpha)$ is to construct all lineage-based positive cell pairs used by LCL and to organize them into mini-batches prior to training, while additionally providing unlabeled cells for the semi-supervised penalty. See Figure 3 of a schematic of positive and negative pairs.

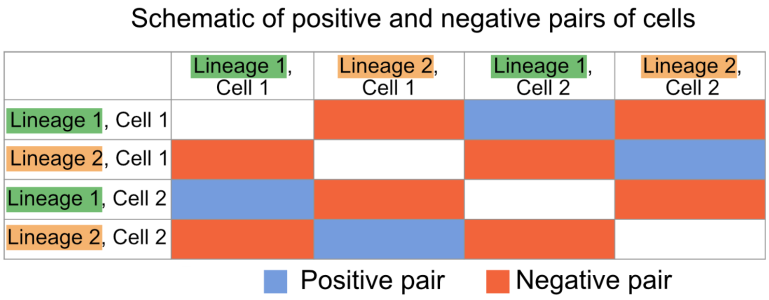


*Figure 3.* Schematic of positive and negative pairs based on 4 cells split across 2 lineages.

For each lineage $L_i$, $\mathcal{C}(\alpha)$ generates $\lceil \alpha \cdot \binom{|L_i|}{2} \rceil$ positive cell pairs. If $\lceil \alpha \cdot \binom{|L_i|}{2} \rceil < |L_i|/2$, we instead generate $\lceil |L_i|/2 \rceil$ pairs to ensure sufficient coverage. We further enforce that every cell appears in at least one positive pair across all generated pairs. Let $S$ denote the total number of positive pairs generated across all lineages.

These $S$ pairs are partitioned into $\lceil S/N \rceil$ mini-batches. Each mini-batch selects $N$ positive pairs from $N$ distinct lineages without replacement whenever possible. If fewer than $N$ lineages remain, the final mini-batch is completed by sampling additional pairs from previously used lineages. The $2N$ cells formed by these $N$ pairs constitute the labeled portion of the batch and are used to compute the supervised contrastive loss.

In addition, for each mini-batch, $\mathcal{C}(\alpha)$ independently samples $U$ cells from an unlabeled dataset (or from held-out cells in the same dataset when no external unlabeled data are provided). These unlabeled cells are not used to form positive or negative pairs. Instead, they are incorporated into the training process through an auxiliary entropy-based penalty that encourages their embeddings to align with the lineage-informed structure learned from the labeled pairs.

#### C.3. Main Algorithm.

After the cell-pair generator has created the batches, they are passed through the main algorithm, and its pseudocode for this process is provided in Algorithm 1.

#### C.4. Details of train/test cells and labeled/unlabeled cells

**Train/validation/test cells (for evaluating generalization).** We first split cells into *training*, *validation*, and *test* sets. The validation set is used only for early stopping during LCL training. After training, we compute embeddings for both training and test cells by running the trained encoder forward on each set (without using test labels to fit either the encoder or downstream predictors). In the "train-test split" setting, the test set contains (approximately) 10% of cells from each lineage, so the set of lineages overlaps between train and test; in the "unseen-lineages" setting, entire lineages are withheld so train/test lineages are disjoint.

**Labeled vs. unlabeled cells (within an LCL mini-batch).** Within each LCL mini-batch, *labeled cells* refer to the $2N$ cells that participate in the lineage-based positive pairs (and hence determine positive vs. negative relationships for the contrastive loss). Separately, when semi-supervision is enabled, we additionally sample $U$ *unlabeled cells* that do not form positive pairs and are not treated as negatives; instead, they contribute only through the auxiliary entropy-based penalty term. In our implementation, unlabeled cells are drawn either from (a) an external unlabeled dataset or (b) a held-out unlabeled pool when no external dataset is provided. In all cases, unlabeled cells are *not* used to define the supervised contrastive labels (i.e., they never create positive pairs).

**Interaction with KNN-based lineage prediction.** For the KNN-based evaluation, the distinction between training and test cells is the only split that directly enters the analysis. Specifically, we train a KNN classifier using the embeddings of

**Algorithm 1** Lineage-aware Contrastive Learning (LCL) with optional semi-supervision

**Input:** Batch lineage count $N \in \mathbb{Z}_+$, size factor $\alpha \in (0, 1]$, temperature $\tau > 0$, penalty weight $\lambda \geq 0$, unlabeled batch size $U \in \mathbb{Z}_{\geq 0}$, cell-pair generator $\mathcal{C}(\alpha)$
Initialize base encoder $f(\cdot)$ and projection head $g(\cdot)$
**for** each batch of $N$ lineage pairs from $\mathcal{C}(\alpha)$ **do**
  **for** $i \in \{1, \dots, N\}$ **do**
    Sample one positive pair $(x_{2i-1}, x_{2i})$ from a lineage $L_i$
    $z_{2i-1} \leftarrow \mathrm{norm}(g(f(x_{2i-1})))$.
    $z_{2i} \leftarrow \mathrm{norm}(g(f(x_{2i})))$
  **end for**
  $Z \leftarrow [z_1, \dots, z_{2N}]$
  **for** Each anchor cell $m \in \{1, \dots, 2N\}$ **do**
    $p(m) \leftarrow$ the index of $m$'s positive partner in $Z$
    $\ell_m \leftarrow -\log \frac{\exp(\mathrm{sim}(z_m, z_{p(m)})/\tau)}{\sum_{k=1}^{2N} \mathbb{1}[k \neq m] \exp(\mathrm{sim}(z_m, z_k)/\tau)}$
  **end for**
  $\mathcal{L}_{\mathrm{con}} \leftarrow \frac{1}{2N} \sum_{m=1}^{2N} \ell_m$
  **if** $\lambda > 0$ **and** $U > 0$ **then**
    Sample unlabeled cells $\{x_u^{(\mathrm{unl})}\}_{u=1}^{U}$
    $u_u \leftarrow \mathrm{norm}(g(f(x_u^{(\mathrm{unl})})))$ for $u \in \{1, \dots, U\}$
    $p_{u,k} \leftarrow \mathrm{softmax}_k(\mathrm{sim}(u_u, z_k)/\tau)$ for $k \in \{1, \dots, 2N\}$
    $\mathcal{L}_{\mathrm{pen}} \leftarrow \frac{1}{U} \sum_{u=1}^{U} \Big( - \sum_{k=1}^{2N} p_{u,k} \log p_{u,k} \Big)$
    $\mathcal{L} \leftarrow \mathcal{L}_{\mathrm{con}} + \lambda \cdot \mathcal{L}_{\mathrm{pen}}$
  **else**
    $\mathcal{L} \leftarrow \mathcal{L}_{\mathrm{con}}$
  **end if**
  Update $f(\cdot), g(\cdot)$ by minimizing $\mathcal{L}$
**end for**
**return** trained $f(\cdot)$ and $g(\cdot)$ (embedding $z = \mathrm{norm}(g(f(x)))$) =0

training-set cells together with their lineage labels, and we then evaluate classification accuracy on the embeddings of the held-out test cells. The labeled/unlabeled distinction applies only during LCL training and influences the learned embedding, but it does not affect how the KNN classifier is trained or evaluated, which is strictly based on the train/test partition.

**Interaction with KL-divergence–based fate composition prediction.** For the KL-divergence evaluation, we again use a train/test split to assess generalization, ensuring that prediction targets are constructed separately within each split to avoid information leakage across time points. For a given split, lineage-level fate compositions are computed using only future-time-point cells from that split (e.g., Day 28 in the CellTag analysis) and are then assigned as targets to early-time-point cells from the same split (e.g., Day 12 in the CellTag analysis). Then, the linear decoder is trained using early-time-point embeddings from the training set with composition targets derived exclusively from training-set future cells, and the KL divergence is evaluated on the test set using targets derived exclusively from test-set future cells. Notably, we fit the linear decoder on only the training split, using an internal training–validation split for early stopping.

### D. Data analysis details

In Section D.1, we document the overall preprocessing pipeline prior to using LCL. In Sections D.2, D.3, D.4, and D.5, we provide summary statistics of the LARRY, CellTag, CellTag-multi, and melanoma sequential-treatment datasets, respectively, and also document the hyperparameters we use for each dataset's analyses. In Section D.6, we document how we simulated pseudo-real data. In Sections D.7 and D.9, we provide more details on the CH Index and compositional analysis, respectively. In Section D.10, we document the full procedure on how we computed the integrated gradients. In Section D.11, we provide the details of how we apply other (existing) methods to investigate fate dynamics. In Section D.12, we document how we perform the batch correction when analyzing the CellTag-multi and CellTag dataset together.

#### D.1. General workflow.

We use Scanpy (Wolf et al., 2018) for preprocessing the input data for the LCL model. Initially, we remove cells that lack a lineage barcode and cells that belong to lineages with fewer than five cells. Afterward, we log-normalize the counts and select the top 2000 highly variable genes (HVGs) for further analysis.

#### D.2. Additional details about the LARRY dataset

**Biological insights.** The authors used isolated unipotent (i.e., one future fate) and pluripotent (i.e., multiple future fates) progenitor cells, using single-cell RNA sequencing to track their fate progression over six days (Weinreb et al., 2020). The primary objective is to determine whether HSPCs follow a continuum of states or discrete hierarchical paths during differentiation. Their findings revealed that progenitors form a structured continuum, and specific progenitors belonged to lineages that resulted in cells differentiating into either a single cell type at a later time (i.e., unipotent) or multiple cell types (i.e., pluripotent).

Table 1 and 2 show the summary of cell types and time points. Specifically, the abbreviations of cell types are basophils ("Baso"), mast cells ("Mast"), megakaryocytes ("Meg"), eosinophils ("Eos"), migratory dendritic cells with marker Ccr7+ ("Ccr7_DC"), and plasmacytoid dendritic cells ("pDC"). Figure 4B shows the number of cells per lineage.

*Table 1.* Cell Type Distribution in the LARRY Dataset

| Cell Type | Count | Proportion |
|---|---|---|
| Undifferentiated | 18426 | 0.448 |
| Neutrophil | 7555 | 0.184 |
| Monocyte | 7350 | 0.179 |
| Baso | 5075 | 0.124 |
| Mast | 1249 | 0.030 |
| Meg | 828 | 0.020 |
| Erythroid | 315 | 0.008 |
| Eos | 147 | 0.004 |
| Lymphoid | 78 | 0.002 |
| Ccr7_DC | 39 | 0.001 |
| pDC | 31 | 0.001 |

*Table 2.* Time Point Distribution in the LARRY Dataset

| Time Point | Count | Proportion |
|---|---|---|
| Day 6 | 27081 | 0.66 |
| Day 4 | 12266 | 0.30 |
| Day 2 | 1746 | 0.04 |

**Training details.** The hyperparameters we use to train the LCL model with the full LARRY dataset (as shown in Figure 6 in the main text) are $\{N = 270, \alpha = 0.4, \tau = 0.5\}$. When using the LARRY dataset to evaluate lineage predictions, the test set includes 10% of cells from each lineage (which we call the "train-test split"), and we use $\{N = 270, \alpha = 0.5, \tau = 0.5\}$. Additionally, when using the LARRY dataset to evaluate the prediction of lineages where the test set includes all cells from 10% of the lineages (which we call the "unseen lineages split"), we use $\{N = 250, \alpha = 0.45, \tau = 0.5\}$.

During our analysis of the LARRY dataset, we observed many lineages with very few cells (see Figure 4B). We observe that these lineages with few cells often deteriorated LCL's performance because there are not enough cell pairs for our contrastive loss function. Hence, we also experimented with two other variants of the LARRY dataset, in which we subset the cells to only the largest lineages. For example, when training only on the cells in the 200 largest lineages from the LARRY dataset with a train-test split, the hyperparameters are $\{N = 150, \alpha = 0.8, \tau = 0.5\}$, while for the unseen lineages split, we use $\{N = 140, \alpha = 0.8, \tau = 0.5\}$. Table 11 reports the number of cells in each subset of the LARRY dataset.

We change the batch size $N$ when analyzing different cell subsets in the LARRY data because the number of available lineages cannot be smaller than $N$. The differences in batch size $N$ and size factor $\alpha$ for the same subset of cells under different experimental conditions are due to system RAM limitations. For scVI and Supervised UMAP, we use the default settings.

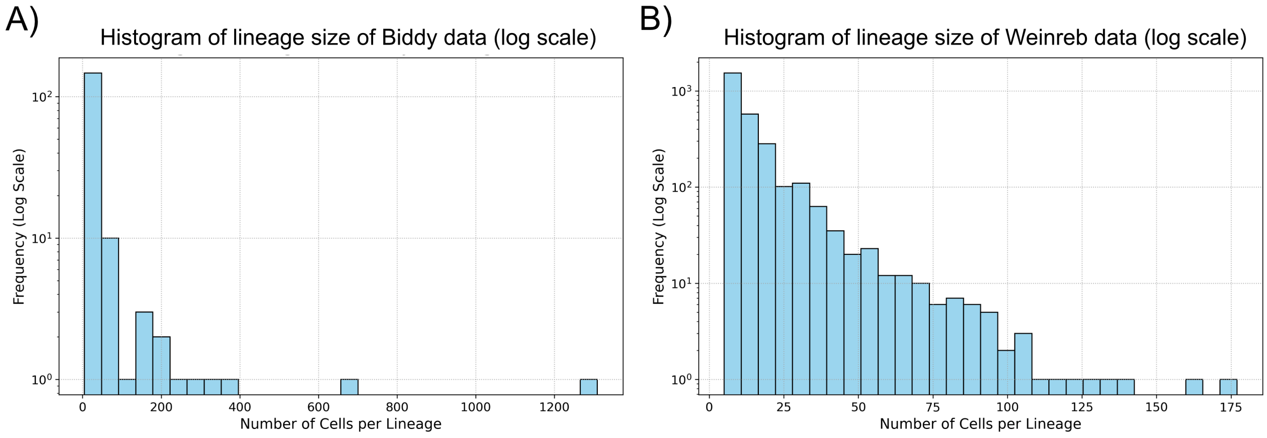


*Figure 4.* **A)** Number of cells per lineage in the CellTag dataset. **B)** Number of cells per lineage in the LARRY dataset.

#### D.3. Additional details about the CellTag dataset.

**Biological insights.** CellTagging, a sequential round of lentiviral barcoding followed by single-cell RNA sequencing, allows the authors to track both the lineage history and identity of individual cells throughout the 28-day reprogramming process (Biddy et al., 2018). See Figure 5 for a visualization of the dataset. The authors aimed to understand cellular heterogeneity and the distinct lineage trajectories that emerge during direct reprogramming, and to identify molecular markers associated with successful lineage conversion. The study revealed two main trajectories: one leading to successful reprogramming into iEPs and another resulting in a “dead-end" state, with early gene expression patterns of specific genes predicting which path cells would take.

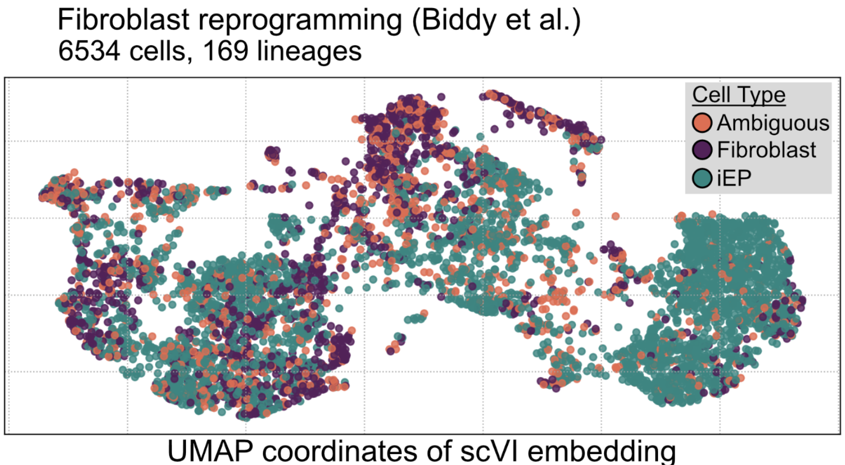


*Figure 5.* UMAP of the CellTag data, where cells are colored by cell type.

Table 3 and 4 summarize the cell types and time points. Figure 4A shows the number of cells per lineage.

*Table 3.* Cell Type Distribution in the CellTag Dataset

| Cell Type | Count | Proportion |
|---|---|---|
| iEP | 3669 | 0.56 |
| Fibroblast | 1508 | 0.23 |
| Ambiguous | 1357 | 0.21 |

*Table 4.* Time Point Distribution in the CellTag Dataset

| Time Point | Count | Proportion |
|---|---|---|
| Day 28 | 2451 | 0.375 |
| Day 21 | 1940 | 0.297 |
| Day 15 | 942 | 0.144 |
| Day 12 | 967 | 0.148 |
| Day 9 | 213 | 0.033 |
| Day 6 | 21 | 0.003 |

**Training details.** The hyperparameters we use to train the LCL model with the CellTag dataset are $\{N = 50, \alpha = 0.04, \tau = 0.5\}$ in all experiments. For scVI and Supervised UMAP, we use the default settings.

#### D.4. Additional details about the CellTag-multi dataset.

**Biological insights.** CellTag-multi extends CellTagging by enabling recovery of heritable lentiviral CellTag barcodes in both scRNA-seq and scATAC-seq, allowing clonal tracking of transcriptional and epigenomic states within the same biological system (Jindal et al., 2024). In the fibroblast-to-iEP direct reprogramming setting, the authors used this multi-omic lineage readout to further link the early cellular state to the eventual reprogramming outcome, beyond what is previously reported in the literature (Biddy et al., 2018). This dissects fate-specific gene-regulatory programs that distinguish on-target iEP trajectories from off-target/dead-end fates.

Tables 5 and 6 summarize the cell types and time points. Here, 'Transition' denotes states in between MEFs and reprogrammed/dead-end identities, while 'Early' denotes early transition reprogramming/transition state shortly after induction. We note that the original data provided additional subtypes for each of these major cell types, which is why Figure 7C in the main text shows different shades for each cell type. We also note that while there are explicitly labeled "Dead-end" cells in the CellTag-multi dataset, these correspond to "Fibroblast" cells in the CellTag dataset that are sequenced at later time points (i.e., dead-end, in the sense that the cells failed to reprogram into iEPs).

*Table 5.* Major Cell Type Distribution in CellTagMulti Dataset

| Major Cell Type | Count | Proportion |
|---|---|---|
| Transition | 9596 | 0.432 |
| Early | 7330 | 0.330 |
| Fibroblast | 2446 | 0.110 |
| iEP | 1761 | 0.079 |
| Dead-end | 1105 | 0.050 |

*Table 6.* Time Point Distribution in the CellTag-multi Dataset

| Time Point | Count | Proportion |
|---|---|---|
| Day 3 | 10890 | 0.49 |
| Day 12 | 3258 | 0.15 |
| Day 21 | 8090 | 0.36 |

**Training details.** The specific details of our analysis integrated CellTag-multi and CellTag are located in Appendix E.6.

#### D.5. Additional details about the melanoma sequential-treatment dataset.

**Biological setting.** To evaluate LCL in a biological system distinct from hematopoietic differentiation and fibroblast reprogramming, we analyzed a melanoma clonal-tracing dataset from Schaff et al. (Schaff et al., 2024). This study combined single-cell RNA sequencing with lineage barcoding to track melanoma cell clones through prolonged and sequential drug treatments. Cells are first exposed to one treatment condition, including cisplatin (Cis), cobalt chloride ($CoCl_2$), or dabrafenib plus trametinib (Dab/Tram), and surviving clones are then subjected to a second-stage treatment. Individual clones exhibited heterogeneous downstream outcomes, including growth, survival, or death, under different treatment sequences. This dataset therefore provides a complementary setting for assessing LCL in a cancer system shaped by treatment-induced selective pressure rather than developmental differentiation.

**Analysis setup.** We apply LCL to cells sampled after the first-stage treatment, using clone identities as lineage labels during contrastive training. We evaluate the learned embedding using two downstream tasks: KNN lineage prediction against a random baseline, and KL-divergence prediction of future treatment-response composition against a naive baseline. For the compositional analysis, we defined each lineage's future outcome by the composition of its surviving descendants across second-stage treatment conditions. This yields a treatment-response composition vector for each clone. We then train a linear decoder on first-stage LCL embeddings to predict these clone-level future survivor compositions and evaluate its performance using KL divergence. The quantitative results are reported in Appendix E.5.

#### D.6. Simulation details on pseudo-real data.

We report additional details on how we simulated the pseudo-real scLT dataset. To generate simulated lineages, genes are first ranked by variance across all cells. For a specific difficulty level $\beta$, we select the top $\beta \cdot p$ genes and reduce the dimensionality to $k = \max\{50, \lceil \beta \cdot p \rceil\}$ principal components. Leiden clustering with default parameters is then applied to the resulting $k$ PCs to assign ground-truth lineage labels.

We report the Leiden clustering resolution for various difficulty levels $\beta$ in Table 7. Recall that a larger $\beta$ represents a dataset that is "easier," whereby there is more information to predict a cell's lineage from its gene expression profile. We change the cluster resolution across different values of $\beta$ to accentuate the differences across $\beta$ better. We also perform a separate experiment, setting the cluster resolution to 45 for all values of $\beta$. In that experiment, we see the same trend as in Figure 5 in the main text – the CH Index is magnitudes higher with LCL than with Supervised UMAP, which is magnitudes higher than that with scVI. However, because the differences between the two methods (with $\beta$ fixed) are much larger than the differences between $\beta$'s (for the same method), we choose to vary the cluster resolution instead to accentuate both the differences across methods and $\beta$'s at the same time.

*Table 7.* Parameters for generating the pseudo-real datasets

| $\beta$ | **Clustering res.** | **Num. of lineages** |
|---|---|---|
| 0.1 | 65 | 1221 |
| 0.3 | 55 | 970 |
| 0.5 | 45 | 797 |
| 0.7 | 35 | 597 |
| 0.9 | 25 | 385 |

#### D.7. Additional details about the Calinski-Harabasz Index.

Suppose there are $n$ cells assigned to $K$ clusters. The Calinski-Harabasz (CH) Index evaluates clustering by comparing the between-cluster dispersion (BCSS) to the within-cluster dispersion (WCSS), adjusted for the number of clusters and data points,

$$\text{CH Index} = \frac{\text{BCSS}/(K-1)}{\text{WCSS}/(n-K)}.$$

In essence, a higher CH Index indicates better cluster separation and compactness.

Here, the Between-Cluster Sum of Squares (BCSS) captures the separation between clusters by measuring the squared distances between each cluster centroid and the overall data centroid,

$$\text{BCSS} = \sum_{k=1}^{K} n_k \|c_k - c\|_2^2,$$

where $n_k$ denotes the number of cells in cluster $k$, $c_k$ is cluster center for cluster $k$, and $c$ is the global center of the dataset. The Within-Cluster Sum of Squares (WCSS) measures how compact the clusters are by summing the squared distances between points and their cluster centroids,

$$\text{WCSS} = \sum_{k=1}^{K} \sum_{i \in C_k} \|x_i - c_k\|_2^2,$$

where $C_k \subseteq \{1, \ldots, n\}$ is the set of indices in the dataset of $n$ cells that are assigned to cluster $k$, and the $i$th cell has coordinates $x_i$.

#### D.8. Additional details about baseline embedding methods used in benchmarking.

To benchmark LCL, we compare its learned embedding against four alternative representations: scVI, Supervised UMAP, a cross-entropy–trained neural embedding, and a triplet-loss–trained neural embedding. For all methods, we use the resulting low-dimensional representations as inputs to the same downstream evaluation pipelines described above: (i) $K$-nearest-neighbor lineage prediction, and (ii) future cell-type composition prediction using a linear decoder followed by KL-divergence evaluation. Thus, differences in downstream performance reflect the quality of the learned embeddings rather than differences in the evaluation procedure.

**scVI.** We train scVI (Lopez et al., 2018) on the training split using raw count data as input, following the standard `scvi-tools` workflow. For each dataset, counts are stored in a dedicated `counts` layer, and scVI is trained using the `SCVI` model with its default optimization procedure. To ensure consistent feature spaces between training and test cells, highly variable genes are selected using only the training data, and the test data are subset to the same gene set before extracting latent representations. For the benchmarking results reported in the main text, we use a 10-dimensional scVI latent embedding. After fitting scVI on the training data, we extract latent representations for both training and held-out test cells using `model.get_latent_representation()`.

**Supervised UMAP.** We use Supervised UMAP (McInnes et al., 2018) as a nonparametric supervised embedding baseline. The model is fit on the training data using lineage labels as supervision and then applied to the held-out test cells to obtain out-of-sample embeddings. We use `n_neighbors = 15` and a 10-dimensional embedding space, i.e., `n_components = 10`. These embeddings are then passed into the same KNN and KL-divergence evaluation workflows used for LCL.

**Cross-entropy–trained neural embedding.** We also benchmarked against a standard lineage-classification network trained with cross-entropy loss. This baseline tests whether directly predicting lineage identities yields an embedding that is useful for downstream fate-related analyses.

The model consists of a multilayer perceptron encoder followed by a linear classification head. Given an input gene-expression vector $x_i \in \mathbb{R}^p$, the encoder maps $x_i$ to a 32-dimensional embedding $h_i \in \mathbb{R}^{32}$:

$$h_i = \phi_{\text{CE}}(x_i).$$

The encoder architecture is

$$p \to 1024 \to 256 \to 32,$$

with batch normalization and ReLU activation after each hidden layer and after the final embedding layer. A linear classifier then maps the embedding to logits over the set of lineage labels:

$$o_i = W h_i + b,$$

where $o_i \in \mathbb{R}^C$ and $C$ denotes the number of lineage classes present in the training split.

The model is trained using the standard multiclass cross-entropy objective,

$$\mathcal{L}_{\text{CE}} = -\frac{1}{n} \sum_{i=1}^{n} \log \left( \frac{\exp(o_{i,y_i})}{\sum_{c=1}^{C} \exp(o_{i,c})} \right),$$

where $y_i$ is the lineage label for cell $i$. Only training-set lineage labels are used to define the classification targets.

We train this model for 50 epochs using AdamW with learning rate $3 \times 10^{-4}$, weight decay $10^{-6}$, and mini-batch size 256. After training, we discard the final classification layer and use the 32-dimensional hidden representation $h_i$ as the embedding for both training and test cells. These embeddings are evaluated using the same KNN and KL-divergence procedures used for LCL.

**Triplet-loss neural embedding.** We further compare LCL with a metric-learning baseline trained using triplet margin loss. This baseline directly encourages cells from the same lineage to lie closer together in embedding space than cells from different lineages, but unlike LCL, it relies on sampled anchor-positive-negative triplets rather than LCL's batchwise contrastive objective.

For each anchor cell $x_a$ with lineage label $y_a$, we randomly sampled: (i) a positive cell $x_p$ from the same lineage, satisfying $y_p = y_a$, and (ii) a negative cell $x_n$ from a different lineage, satisfying $y_n \neq y_a$. The model encoder maps each input to a 32-dimensional embedding,

$$z_a = \phi_{\text{tri}}(x_a), \qquad z_p = \phi_{\text{tri}}(x_p), \qquad z_n = \phi_{\text{tri}}(x_n),$$

which is then $\ell_2$-normalized before computing the loss.

The encoder architecture is

$$p \to 1024 \to 256 \to 32,$$

with batch normalization and ReLU activation after the first two hidden layers. Unlike the cross-entropy baseline, no classification head is used. The final embedding is normalized as

$$\tilde{z} = \frac{z}{\|z\|_2}.$$

We optimize the standard triplet margin loss,

$$\mathcal{L}_{\text{triplet}} = \frac{1}{B}\sum_{i=1}^{B} \max\left\{d(\tilde{z}_{a_i}, \tilde{z}_{p_i}) - d(\tilde{z}_{a_i}, \tilde{z}_{n_i}) + m,\, 0\right\},$$

where $B$ is the mini-batch size, $d(\cdot,\cdot)$ is the Euclidean distance, and the margin is set to $m = 1.0$.

We train the triplet-loss model for 50 epochs using AdamW with learning rate $3 \times 10^{-4}$, weight decay $10^{-6}$, and mini-batch size 256. After training, the 32-dimensional normalized embeddings are extracted for both training and held-out test cells and evaluated using the same KNN and KL-divergence benchmarks as the other methods.

#### D.9. Additional details about the compositional analysis.

In the LARRY dataset, we designate Day 2 as the early time point, with all cells labeled Undifferentiated, and Day 6 as the future time point. We focus on only the Day 2 cells in the Monocyte-Neutrophil trajectory, as defined by the original authors (Weinreb et al., 2020). By Day 6, the cell types diversify, and the three largest cell type populations are neutrophils, monocytes, and undifferentiated cells. We compute our compositional vector among these three cell types on Day 6.

In the CellTag dataset, we designate Day 12 as the early time point and Day 28 as the future time point. Both time points include three cell types: iEP, fibroblast, and ambiguous. However, by Day 28, the iEP population are notably larger proportionally, making up 72% of the total cells, compared to 53% at Day 12. We compute our compositional vector among these three cell types on Day 28.

#### D.10. Additional details about our usage of integrated gradients.

To identify genes that most strongly contribute to lineage-resolved structure learned by LCL, we apply Integrated Gradients (IG) (Sundararajan et al., 2017) to a trained LCL model in a post hoc manner. Rather than attributing individual embedding coordinates, we define a *lineage-specific scalar objective* measuring how strongly a cell aligns with its lineage centroid in the learned embedding space.

**Embedding and lineage centroids.** Let $x \in \mathbb{R}^p$ denote the (log-normalized) gene-expression vector for a cell across $p$ genes. LCL consists of a base encoder $f(\cdot)$ and projection head $g(\cdot)$, producing a normalized embedding

$$z(x) \;=\; \mathrm{norm}(g(f(x))) \in \mathbb{R}^d, \qquad \mathrm{norm}(u) \;=\; \frac{u}{\|u\|_2 + \varepsilon},$$

with a small $\varepsilon > 0$ for numerical stability. For each lineage $\ell$, let $\mathcal{I}_\ell$ be the set of indices of cells in that lineage. We compute the lineage centroid in embedding space as,

$$\mu_\ell \;=\; \mathrm{norm}\left(\frac{1}{|\mathcal{I}_\ell|}\sum_{i\in\mathcal{I}_\ell} z(x_i)\right).$$

In practice, embeddings $z(x_i)$ are computed for all cells without gradients; centroids are computed for the subset of lineage considered in the IG analysis.

**Lineage-specific scalar attribution target.** For a fixed lineage $\ell$, we define a scalar function

$$F_\ell(x) \;=\; \cos\big(z(x), \mu_\ell\big).$$

Intuitively, $F_\ell(x)$ measures how aligned a cell's embedding is with the lineage's centroid direction.

**Integrated Gradients.** Given a baseline input $x'$ (we use $x' = 0 \in \mathbb{R}^p$, i.e. all genes set to zero; alternatively one may use the global mean expression), the IG attribution for gene $j \in \{1, \ldots, p\}$ is

$$\mathrm{IG}_j(x; x', F_\ell) \;=\; (x_j - x'_j)\int_0^1 \frac{\partial F_\ell\big(x' + \alpha(x - x')\big)}{\partial x_j}\, d\alpha.$$

We approximate the integral using a fixed number of steps (default: $m = 100$) with Gauss–Legendre quadrature (as implemented by the Python package `Captum`).

**Lineage-level and global gene scores.** For each lineage $\ell$, we randomly sample up to $M = 200$ cells (or all cells if $|\mathcal{I}_\ell| < M$) and compute IG for each sampled cell. We summarize per-lineage gene importance by the mean absolute attribution:

$$s_{\ell,j} \;=\; \frac{1}{|S_\ell|}\sum_{i\in S_\ell} \left|\mathrm{IG}_j(x_i; x', F_\ell)\right|,$$

where $S_\ell \subseteq \mathcal{I}_\ell$ is the set of sampled cell indices for lineage $\ell$. To obtain a global ranking across lineages, we average across the $L$ analyzed lineages:

$$s_j \;=\; \frac{1}{L}\sum_{\ell=1}^{L} s_{\ell,j}.$$

Genes with large $s_j$ are those whose expression consistently has a large influence (in magnitude) on alignment to lineage centroids in the LCL embedding.

#### D.11. Additional details about applying other methods when identifying pathways related to fate commitment.

- **Usage of VariancePartition**: We apply the VariancePartition analysis (Hoffman & Schadt, 2016) based on the `variancePartition` R package (found on Bioconductor), which uses a mixed-effects model to model the amount of variance explained in each gene's expression by each covariate. We use a regression form of `(1 | Lineage_label) + (1 | Cell_type) + (1 | Time_point)` and the `variancePartition::fitExtractVarPartModel` function.
- **Usage of GEMLI**: We apply GEMLI (Eisele et al., 2024), which is modified from `predict_lineages` function in the `GEMLI` R package (found on GitHub, `UPSUTER/GEMLI`). The authors design this function to take the normalized single-cell gene expression matrix by default and to compute "marker genes," which are the only features used in the remainder of the function when predicting whether two cells are from the same lineage. However, we code a modification of `predict_lineages` that allows us to avoid computing marker genes (and instead use all the provided features), since we want to apply GEMLI to LCL or scVI coordinates. Otherwise, we use the default GEMLI parameters. Specifically, we use `fast=TRUE`, `repetitions=100`, and `sample_size=2/3`. We set `desired_cluster_size=c(50,200)` since our application of GEMLI on the CellTag-multi dataset focuses on only the cells in the largest 50 lineages.

  Then, to predict, we use the `test_lineages` function with `valid_fam_sizes=c(50,200)` to compute the true and false positive rates.

- **Usage of CoSpar**: We apply CoSpar (Wang et al., 2022), which is the `cospar` Python package (found on GitHub at `AllonKleinLab/cospar`). We use the recommended parameters when using CoSpar on the CellTag-multi data. Specifically, we use `cospar.tmap.infer_Tmap_from_multitime_clones` with the following parameters:
  - `clonal_time_points=["3", "12", "21"]`,
  - `later_time_point="21"`,
  - `smooth_array=[15, 10, 5]`,
  - `sparsity_threshold=0.2`,
  - `intraclone_threshold=0.2`.

- **Usage of GSEA**: We perform GSEA using the `clusterProfiler` R package (Yu et al., 2012). Specifically, we use the `clusterProfiler::gseGO` function using the recommended parameters in their tutorial, `minGSSize=10` and `maxGSSize=500`. We only analyze the biological process ontology (i.e., `ont="BP"`) using the pathways from the `org.Mm.eg.db` (for the mouse).

  The inputs to GSEA are: 1) the integrated gradient of each gene (based on our description in the Section 3.4 in the main text), 2) the correlation between CoSpar's fate bias for differentiating into either reprogramming/dead-end cell types and a normalized gene's expression across those same cells, and 3) the VariancePartition's variance explained among all the genes for the lineage label.

#### D.12. Additional details on the CellTag and CellTag-multi integration analysis

When analyzing the integration between the CellTag and CellTag-multi datasets, we first perform batch correction using Seurat's anchor-based integration framework (Stuart et al., 2019; Hao et al., 2021) prior to applying LCL. This preprocessing step is necessary because the LCL architecture does not explicitly incorporate a batch-effect correction term, as such corrections are not naturally compatible with the contrastive learning objective.

To do this batch correction, we use Seurat's anchoring method. Our empirical experiment demonstrates that this batch correction method is more suitable than using scANVI. The integration features are selected using `Seurat::SelectIntegrationFeatures()` (`nfeatures = 3000`), and cross-dataset anchors are identified using `Seurat::FindIntegrationAnchors()` (default CCA-based integration with PCA dimensions 1-30). The integrated expression assay is then constructed using `Seurat::IntegrateData()` (default `new.assay.name = "integrated"`).

### E. Additional results

In Section E.1, we provide additional results using VariancePartition. In Section E.2, we provide additional results using LCL, scVI, or supervised UMAP for the analyses on either the LARRY and CellTag datasets. In Section E.3, we provide additional GEMLI and GSEA evaluations on the full LARRY dataset. In Section E.4, we provide additional results about the KNN classifier or compositional analysis for both the LARRY and CellTag datasets. In Section E.5, we report additional results for the melanoma sequential-treatment dataset. In Section E.6, we provide additional results on the CellTag-multi and CellTag integrated analysis. In Section E.7, we document the runtime for each analysis, provide guidance on how the hyperparameters impact the runtime, and report the hyperparameters used for each analysis.

#### E.1. Additional results about the VariancePartition analysis.

In the main text, we use VariancePartition (Hoffman & Schadt, 2016) to analyze how much of each gene's expression can be explained by each covariate (lineage label, time point, and cell type). We demonstrate this on the LARRY dataset (Weinreb et al., 2020) in the main text (Figure 3). Here, we display the VariancePartition results for the CellTag and CellTag-multi dataset (Figure 6). We can see that the amount of variance explained by lineage is: (1) highly variable from gene to gene, and (2), on average, is relatively small when compared to cell type and time points. These results further emphasize why learning fate-commitment dynamics from scLT data is difficult – the signal from the lineage labels is not the dominant source of biological information. Unlike other machine learning tasks, where the "largest source of signal" is typically the primary signal of interest, we are instead interested in the biological relevance of secondary signals.

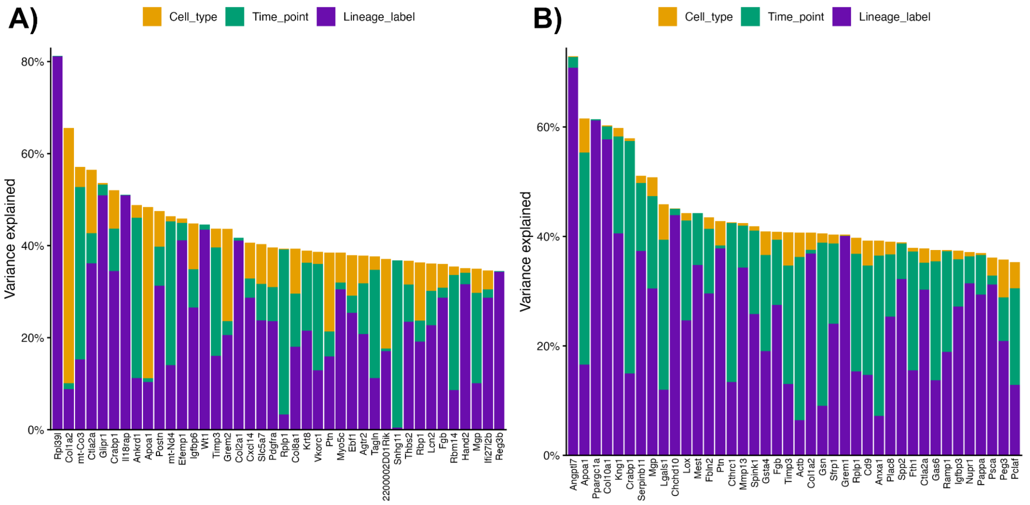


*Figure 6.* The VariancePartition results for CellTag (**A**) and CellTag-multi (**B**), showing the top 40 genes in each respective dataset. The plot is organized similarly to Figure 3 in the main text.

### E.2. Additional results of LARRY and CellTag datasets

Figure 7 shows the LCL embedding for the CellTag dataset. This mirrors Figure 6 in the main text.

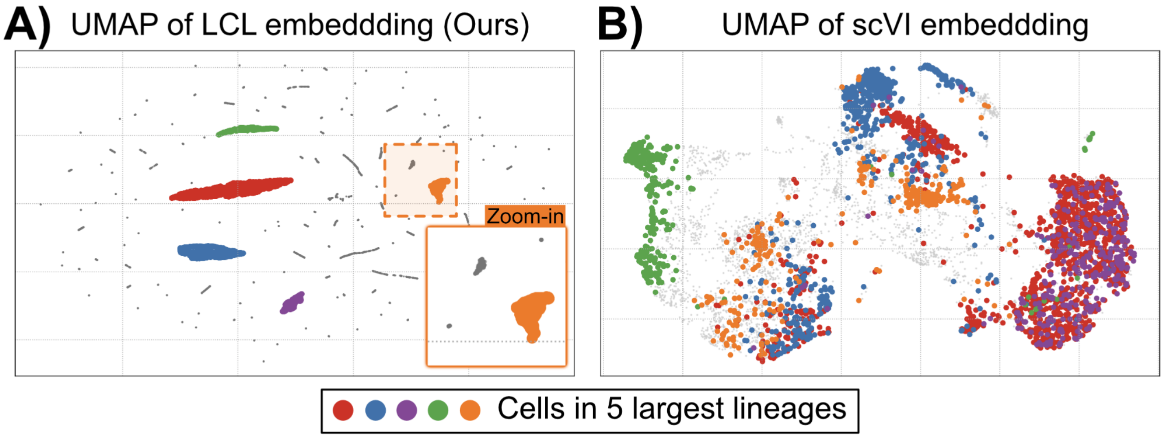


*Figure 7.* UMAP of the LCL embedding (**A**) or the scVI embedding (**B**) for the CellTag dataset. The cells in the largest five lineages are highlighted with a different color in their respective datasets.

Figure 8 shows the Supervised UMAP of both the CellTag and LARRY datasets. These plots contrast the embeddings of LCL and scVI in Figure 6 in the main text. We note that Figure 6 in the main text shows the LCL embedding specifically for the “LARRY” (i.e., with all 41,093 cells) dataset.

Figure 9 shows the UMAP of the LCL embedding of “LARRY (Largest 200)” The embedding also looks “island-like” in the UMAP, illustrating that LCL has found a way to separate many lineages and to cluster cells within the same lineages, even if the lineage contains cells of many different cell types collected at different time points.

### E.3. Additional GEMLI and GSEA evaluation on the full LARRY dataset

The full LARRY dataset is highly heterogeneous, tracking 41,093 cells across 2,813 distinct lineages in a complex hematopoietic differentiation system. To further evaluate whether LCL learns a representation that preserves lineage structure in this heterogeneous setting, we apply GEMLI (Eisele et al., 2024) to compare whether different embeddings can predict if two cells originate from the same lineage. Using the LCL embedding substantially improved GEMLI’s predictive

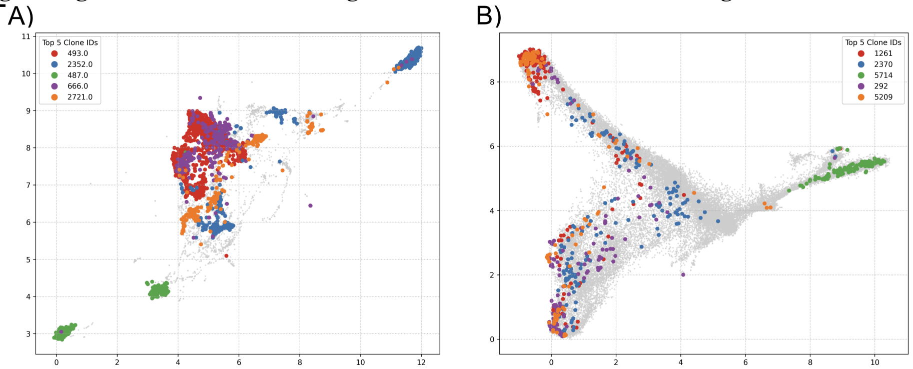


*Figure 8.* **A)** Supervised UMAP of the CellTag dataset. **B)** Supervised UMAP of the LARRY dataset. The cells in the five largest lineages are highlighted in different colors. These colors are the same as those shown in Figure 6 in the main text.

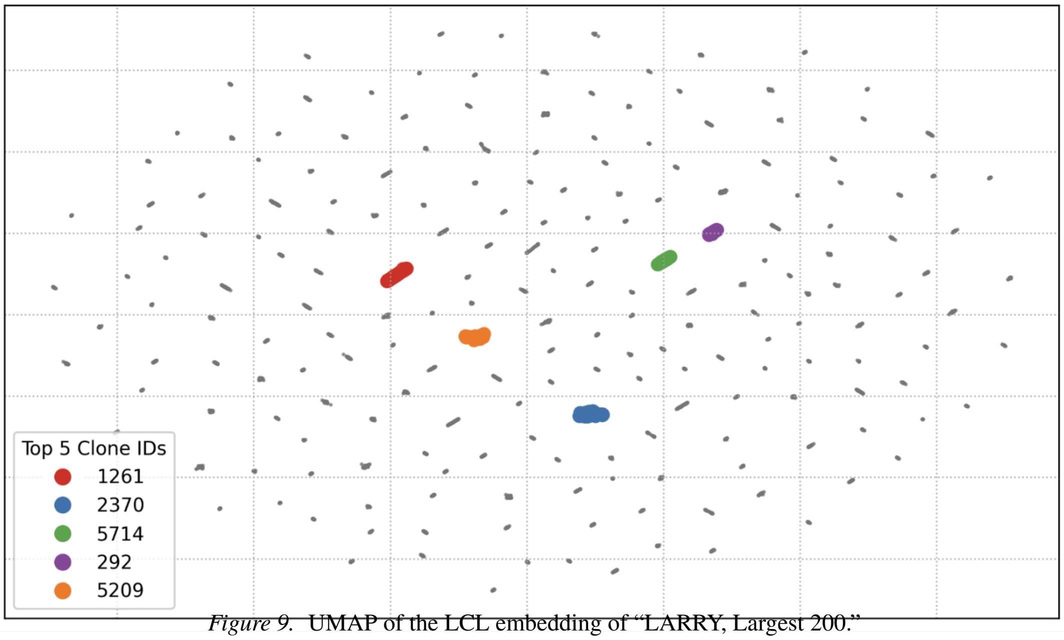


*Figure 9.* UMAP of the LCL embedding of "LARRY, Largest 200."

performance, achieving an AUROC of 0.73, compared to 0.13 using the scVI embedding and 0.16 using log-normalized gene expression features (Figure 10).

We further evaluate whether the LCL embedding supports pathway-level discovery of lineage-associated biological signals. We apply Hotspot-based gene scoring followed by GSEA to compare pathway enrichment across methods. LCL uniquely identified statistically significant fate-associated pathways, including *response to external stimulus* and *oxygen-containing compound*, that are not detected using scVI or CoSpar (Figure 11). These results support the conclusion that LCL improves both lineage-structure recovery and downstream biological discovery on the full LARRY dataset.

#### E.4. Additional results of KNN and compositional analysis

**Evaluating lineage clusters and predicting lineage.** Table 8 shows the CH Index for evaluating if a lineage is clustered together in the embedding and the KNN accuracy for predicting a cell's lineage using the KNN classifier with five nearest neighbors ("KNN(5)"). We show these metrics on both the training and test sets. These correspond to the results in Table 1 in the main text. Specifically, the results in Table 1 correspond to "CellTag" and "LARRY, Largest 200" in Table 8. Recall that a higher CH Index and a higher KNN accuracy indicate that the embedding generalizes better. We bold the highest training and testing values for each dataset in Table 8.

**Generalizing to unseen lineages in the LARRY and CellTag datasets.** We also consider a more challenging approach to constructing a test set, in which the test set cells belong to lineages not present in the training set (i.e., "unseen"). Since the lineages in the training and test sets are mutually exclusive, this experiment assesses a more challenging generalization.

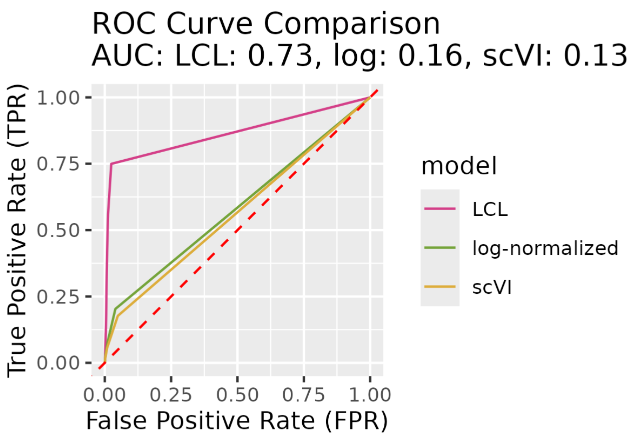


*Figure 10.* ROC curves for GEMLI lineage-pair prediction on the full LARRY dataset using different input representations. GEMLI predicts whether two cells originate from the same lineage using a random forest classifier. LCL achieves an AUROC of 0.73, outperforming scVI and log-normalized gene-expression inputs.

*Table 8.* CH Index and KNN classifier's accuracy for predicting lineage. The values in the "KNN(5) Test" match the values in Table 1 in the main text (since Table 1 shows error, and this table shows accuracy)

| **Dataset** | **Method** | **CH Index** | | **KNN(5)** | |
|---|---|---|---|---|---|
| | | **Train** | **Test** | **Train** | **Test** |
| CellTag | LCL | **15351.3** | **14.7** | **98.98%** | **54.06%** |
| | supUMAP | 66.1 | 11.6 | 71.8% | 37.2% |
| | scVI | 18.1 | 4.9 | 49.7% | 47.1% |
| LARRY, Largest 200 | LCL | **56335.9** | **9.3** | **99.0%** | **13.6%** |
| | supUMAP | 125.4 | 6.2 | 33.7% | 5.9% |
| | scVI | 30.4 | 4.8 | 12.6% | 10.8% |
| LARRY | LCL | **430.1** | **7.6** | **47.9%** | 2.5% |
| | supUMAP | 35.9 | 5.3 | 3.9% | 1.6% |
| | scVI | 12.2 | 3.2 | 4.3% | **3.3%** |

We use a different evaluation metric for this setting. Based on the fitted LCL model, we construct a KNN graph of the test cell's embedding and count the number of test-cell nearest neighbors that are from the same lineage. We are interested in this experiment since, if successful, LCL would be applied to biological systems where *in vitro* lineage barcoding is not possible. Our results demonstrate that LCL offers a modest improvement over scVI and a more substantial improvement over Supervised UMAP.

Table 9 shows the performance of each of the three embedding methods in the "unseen lineages" scenario whereby 10% of the lineages are entirely withheld from the training set, and the cells of these withheld lineages formed the test set. This setting is more difficult because the lineages in the training and test sets do not overlap.

Since the lineages between the training and test sets do not overlap, we use a different evaluation metric. Specifically, the values reported in Table 9 refer to the average percentage of nearest-neighbor test cells (among the five nearest neighbors) that share the same lineage as the test cell in question ("KNN(5)"). A higher number here indicates that the embedding generalizes to unseen lineages, with cells in the same test lineage mapped to similar coordinates in the embedding space. We bold the highest values for each dataset in Table 9.

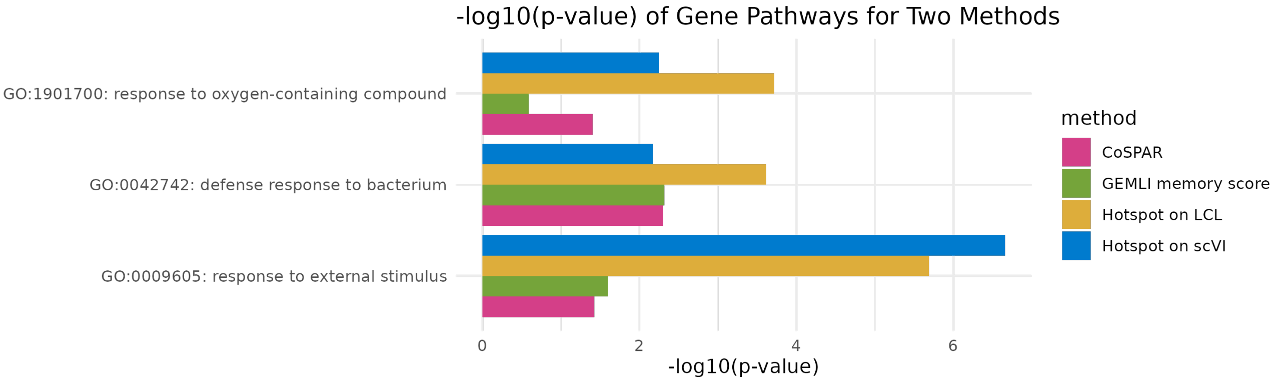


*Figure 11.* Pathway enrichment comparison on the full LARRY dataset using Hotspot-based gene scoring followed by GSEA. The plot compares pathway enrichment across methods using $-\log_{10}(p\text{-value})$. LCL uniquely identifies statistically significant pathways such as response to external stimulus and oxygen-containing compound, which are missed by alternative methods.

*Table 9.* Percentage of nearest-neighbor test cells with the same lineage for unseen lineages

| Dataset | Method | KNN(5) |
|---|---|---|
| CellTag | LCL | 44.90% |
| | supUMAP | 37.20% |
| | scVI | **45.70%** |
| LARRY, Largest 200 | LCL | **50.60%** |
| | supUMAP | 41.40% |
| | scVI | 48.60% |
| LARRY | LCL | **26.90%** |
| | supUMAP | 16.40% |
| | scVI | 21.30% |

### E.5. Additional results on the melanoma sequential-treatment dataset

We evaluate whether LCL generalizes to a cancer system with treatment-driven clonal fate dynamics using the melanoma sequential-treatment dataset from Schaff et al. (Schaff et al., 2024). As summarized in Table 10, LCL outperforms the corresponding baselines for both KNN lineage recovery and KL-divergence prediction of future survivor composition.

*Table 10.* Performance of LCL on the melanoma sequential-treatment dataset. Lower values indicate better performance.

| Evaluation Metric | LCL | Baseline |
|---|---|---|
| KNN Lineage Error | 0.746 | 0.986 |
| KL Divergence for Future Survivor Composition | 1.047 | 1.689 |

The lower KNN lineage error indicates that the LCL embedding captures clone-specific structure in held-out cells, while the reduced KL divergence suggests that first-stage lineage-aware embeddings retain information predictive of future treatment-response compositions. These results extend our evaluation beyond developmental and reprogramming systems, supporting the applicability of LCL to a cancer model with distinct lineage dynamics and sequential treatment perturbations.

### E.6. Additional results on CellTag/CellTag-multi integration

**Batch correction.** Before applying LCL to the experiment assessing whether the lineage labels in CellTag-multi are informative for learning fate dynamics in CellTag, we need to perform batch correction. Figure 12A shows a clear batch effect between the two datasets before any correction. Since LCL does not do a batch correction, Figure 12 shows the resulting UMAP, colored by dataset identity, illustrating an effective batch-correction, standard-anchor-based integration workflow. This integration of gene expression vectors for both datasets provides a common coordinate system on which LCL can be applied to investigate lineage-prediction transfer between related datasets.

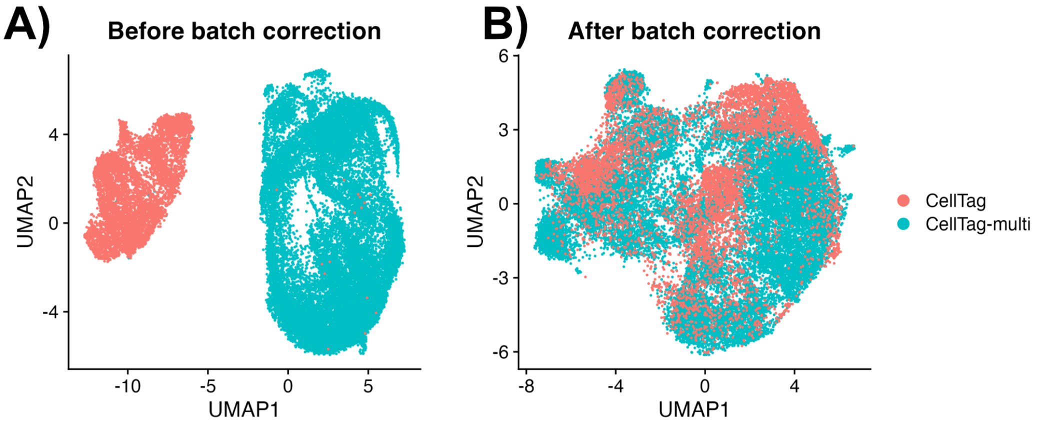


*Figure 12.* **A)** Gene expression of the CellTag and CellTag-multi datasets, where the two datasets are clearly separable due to batch effects. Cells are colored by dataset identity. **B)** Baseline Seurat batch-correction integrating both datasets together.

We note that, even after integrating the two datasets, the lineages in CellTag are not necessarily co-localized. In Figure 13, we see that the largest 5 lineages in CellTag are quite dispersed throughout the integrated gene expression space.

**Size-normalized KNN lineage enrichment (modified KNN).** While the CH Index captures *global* separation of lineage clusters, it may fail to reflect settings in which lineage structure is expressed primarily at a local scale. Moreover, a standard KNN classification accuracy is ill-suited to this integration setting, as it is highly sensitive to lineage-size imbalance and does not provide a meaningful baseline under random mixing. We therefore define a size-normalized neighborhood enrichment metric that directly quantifies whether same-lineage cells are overrepresented in local neighborhoods relative to their global frequency.

Fix a neighborhood size $K$ and let $\mathcal{N}_K(i)$ denote the set of $K$ nearest neighbors of cell $i$ in the embedding space (excluding $i$ itself). For each cell $i$ with lineage label $y_i$, we compute

$$p_{\text{local}}(i) = \frac{1}{K} \sum_{j \in \mathcal{N}_K(i)} \mathbb{1}[y_j = y_i], \qquad p_{\text{global}}(i) = \frac{n_{y_i} - 1}{n - 1},$$

where $n_{y_i}$ is the size of lineage $y_i$ among the $n$ evaluated cells. Here, $p_{\text{global}}(i)$ represents the expected fraction of same-lineage neighbors under random mixing, accounting for lineage size.

We define the per-cell enrichment score as

$$\text{Enrich}(i) = \frac{p_{\text{local}}(i)}{p_{\text{global}}(i)}.$$

An enrichment value greater than 1 indicates that cell $i$'s neighborhood contains more same-lineage cells than expected by chance, while a value near 1 corresponds to no lineage-specific enrichment. We summarize embedding quality using the mean and median enrichment across cells:

$$\text{Enrich}_{\text{mean}} = \frac{1}{n} \sum_{i=1}^{n} \text{Enrich}(i), \qquad \text{Enrich}_{\text{median}} = \text{median}_i \, \text{Enrich}(i).$$

Larger enrichment scores indicate stronger local lineage coherence. We report results with neighborhood sizes of 50 to assess robustness across spatial scales.

##### E.6.1. Integration hyperparameter diagnostics: Calinski–Harabasz and size-normalized KNN enrichment

To summarize integration quality across different semi-supervised LCL settings (e.g., varying $\lambda$ and the number of unlabeled cells $U$), we evaluate each fitted embedding using two complementary diagnostics computed on the CellTag cells (treated as unlabeled during training, but with lineage barcodes available for post hoc evaluation). These metrics capture two distinct

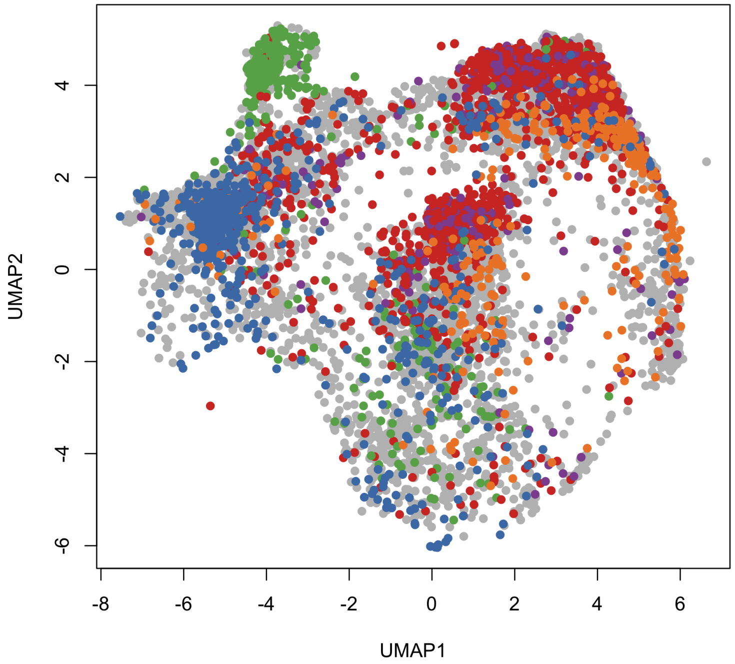


*Figure 13.* The largest 5 lineages of the CellTag dataset in the Seurat batch-corrected embedding space. This plot illustrates the need to run LCL after batch correction, since the true lineages (which LCL will not have access to during training, since CellTag will be treated as an "unlabeled" dataset) are not co-localized in this embedding space.

desiderata: (i) whether lineage structure forms globally separated clusters, and (ii) whether local neighborhoods are enriched for same-lineage cells beyond what would be expected from lineage size alone. As Figure 14 shows, we choose the setting with $\lambda = 0.05$ and $U = 50$ that balances strong local lineage enrichment with moderate global cluster separation, avoiding over-fragmentation while preserving lineage coherence.

#### E.6.2. Marker genes of LCL clusters.

Continuing the analysis of the genes that strongly correlate with fate dynamics (Table 4 in the main text), we first show in Figure 16 the Leiden clustering of the CellTag (unlabeled) cells in the learned LCL embedding. We use the `Seurat::FindClusters` function, with `resolution=0.1`, to learn these 8 clusters. These clusters provide an automated way to separate the large "islands" in the LCL embedding. Afterwards, we use a Wilcoxon test to learn the differentially expressed genes unique to each island using a one-versus-rest strategy used by the `Seurat::FindAllMarkers` function. We filter the list of genes such that: 1) the log fold change of the gene (in the target cluster, relative to all the other cells) is 3 or larger, 2) the percent of cells expressing the gene in the target cluster is at least 30%, and 3) the adjusted p-value is less than 0.05. Only 17 genes remain after this stringent filtering, of which we focus on *Ptn*, *Sfrp1*, and *Shh*, three genes strongly associated with whether a fibroblast cell will successfully reprogram into an iEP.

The genes identified through LCL clustering (*Ptn*, *Sfrp1*, and *Shh*) represent critical molecular divergence points between successful and unsuccessful reprogramming trajectories. *Ptn* (Pleiotrophin) and Sfrp1 (Secreted Frizzled Related Protein 1) are established markers of "dead-end" cells that fail to transition into Induced Endoderm Progenitors (iEPs) (Jindal et al., 2024). *Sfrp1* is a known antagonist of Wnt signaling, and its upregulation is part of a "dead-end" signature that reinforces the fibroblast identity or leads to off-target states. In contrast, the activation of the *Shh* (Sonic Hedgehog) signaling pathway is associated with successful reprogramming into the endoderm lineage. While the CellTag-multi (Jindal et al., 2024) study focuses heavily on the Wnt/TGF-$\beta$ axis to distinguish on-target iEP trajectories from dead-end fates, Shh is a documented driver of definitive endoderm differentiation and embryonic gut development, serving as a hallmark of successful iEP maturation (Guzzetta et al., 2020).

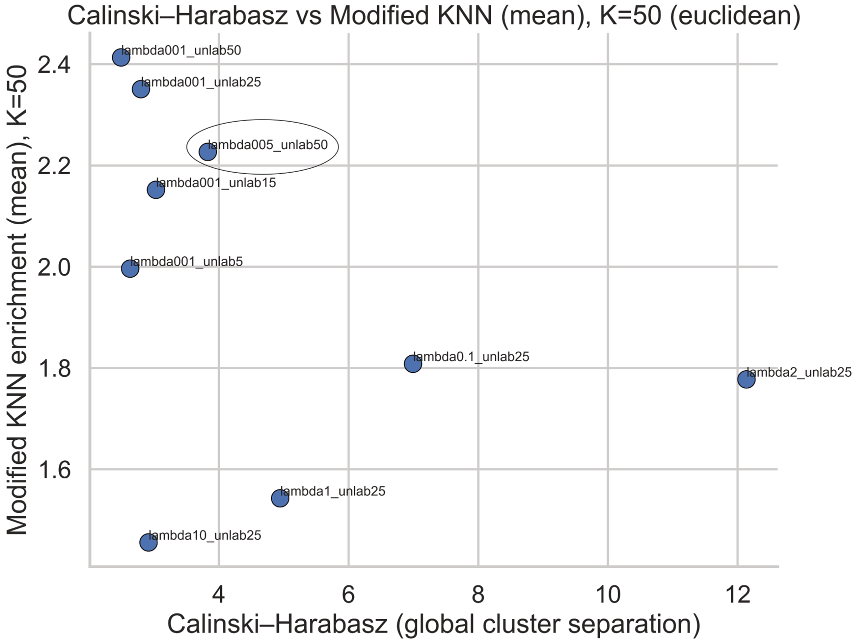


*Figure 14.* Trade-off between global lineage separation (CH Index) and local lineage coherence (size-normalized KNN enrichment, $K = 50$) across LCL integration settings. The circled point denotes the selected hyperparameter setting used in downstream analyses.

We illustrate the co-localization of these key genes in Figure 17, where we contrast the original integrated gene expressions against the LCL embeddings. Specifically, we observe in Figure 17B that genes like *Ptn* and *Sfrp1* co-localize very strongly into certain LCL "islands," which consist of mainly fibroblasts that do not successfully reprogram into iEPs. This reinforces LCL's strength: LCL can isolate the gene expression component that specifically relates to fate dynamics encoded by lineage labels in the CellTag dataset, *even though* LCL did not access or use these lineage labels during training. This demonstrates both LCL's generalizability and potential to advance cell biology.

#### E.7. Additional results on runtime performance

We discuss how LCL scales with different dataset sizes, batch configurations, and hyperparameters. This provides additional insight into the computational feasibility of applying LCL to other large-scale single-cell lineage-tracing datasets. The main factors influencing runtime are the number of cells, mean number of cells per lineage, batch size $N$, size factor $\alpha$, and number of training epochs. Table 11 shows the number of cells, number of cells per lineage, the hyperparameters we use for LCL, and the resulting run time of the LCL fitting procedure.

- **Effect of Number of Cells and Cells per Lineage:** There is a moderate positive correlation between both the number of cells ($r = 0.59$) and the number of cells per lineage ($r = 0.55$) with the runtime. This suggests that datasets with more cells or more cells per lineage tend to require longer runtimes. For instance, the dataset with 41,093 cells and an average of 33 cells per lineage takes 1.78 hours to run.
- **Effect of Batch Size and Positive Pairs:** Larger datasets with more positive pairs take longer to process, especially when the size factor is increased.
- **Impact of Hyperparameters:** The batch size and size factor directly influence runtime, with larger batch sizes reducing the number of total batches and slightly improving GPU utilization. The dataset's number of epochs and complexity (e.g., simulation datasets with varying beta values) also contribute significantly to runtime variability.
- **Efficiency on Google Colab:** Training the model on Google Colab's NVIDIA L4 GPU is efficient for medium-sized datasets (up to 1.7 million pairs), with average run times of less than 2 hours. The 2.3 million trainable parameters and efficient use of GPU and system resources allow LCL to handle various datasets without exceeding resource limits.

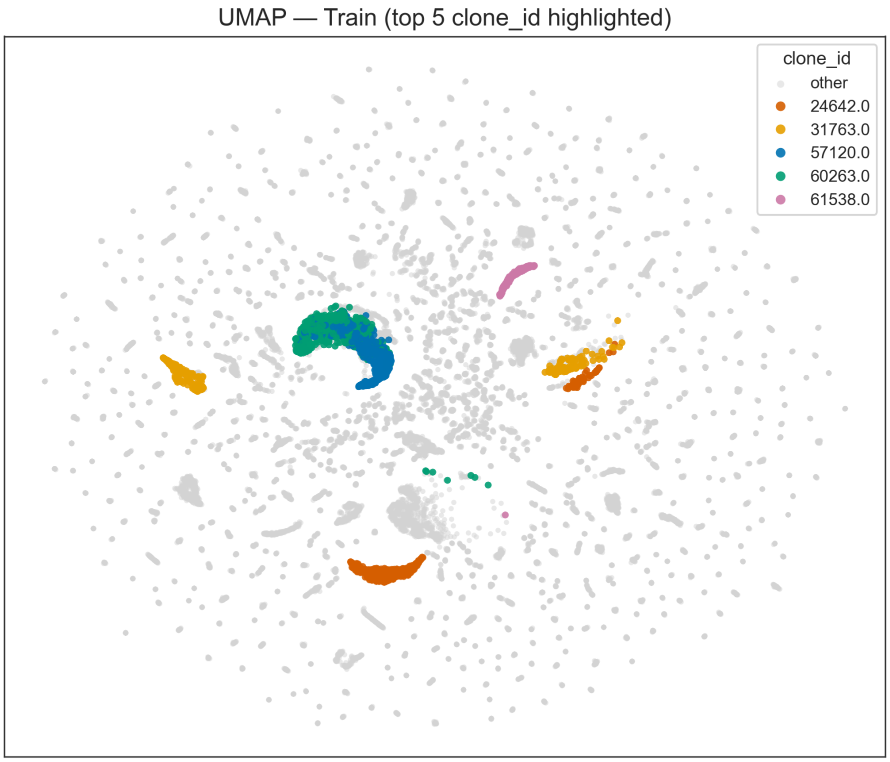


*Figure 15.* UMAP visualization of the LCL embedding for the CellTag-multi training set, with the five largest lineages highlighted. Cells belonging to the top five lineages (by clone size) are colored, while all other cells are shown in gray. This visualization illustrates the within-lineage coherence and spatial separation learned by LCL on the labeled reference dataset.

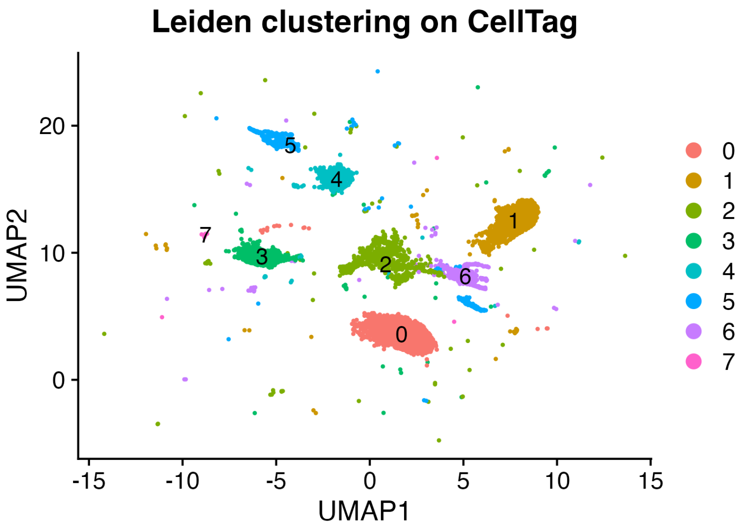


*Figure 16.* Leiden clustering of the CellTag dataset's LCL embedding.

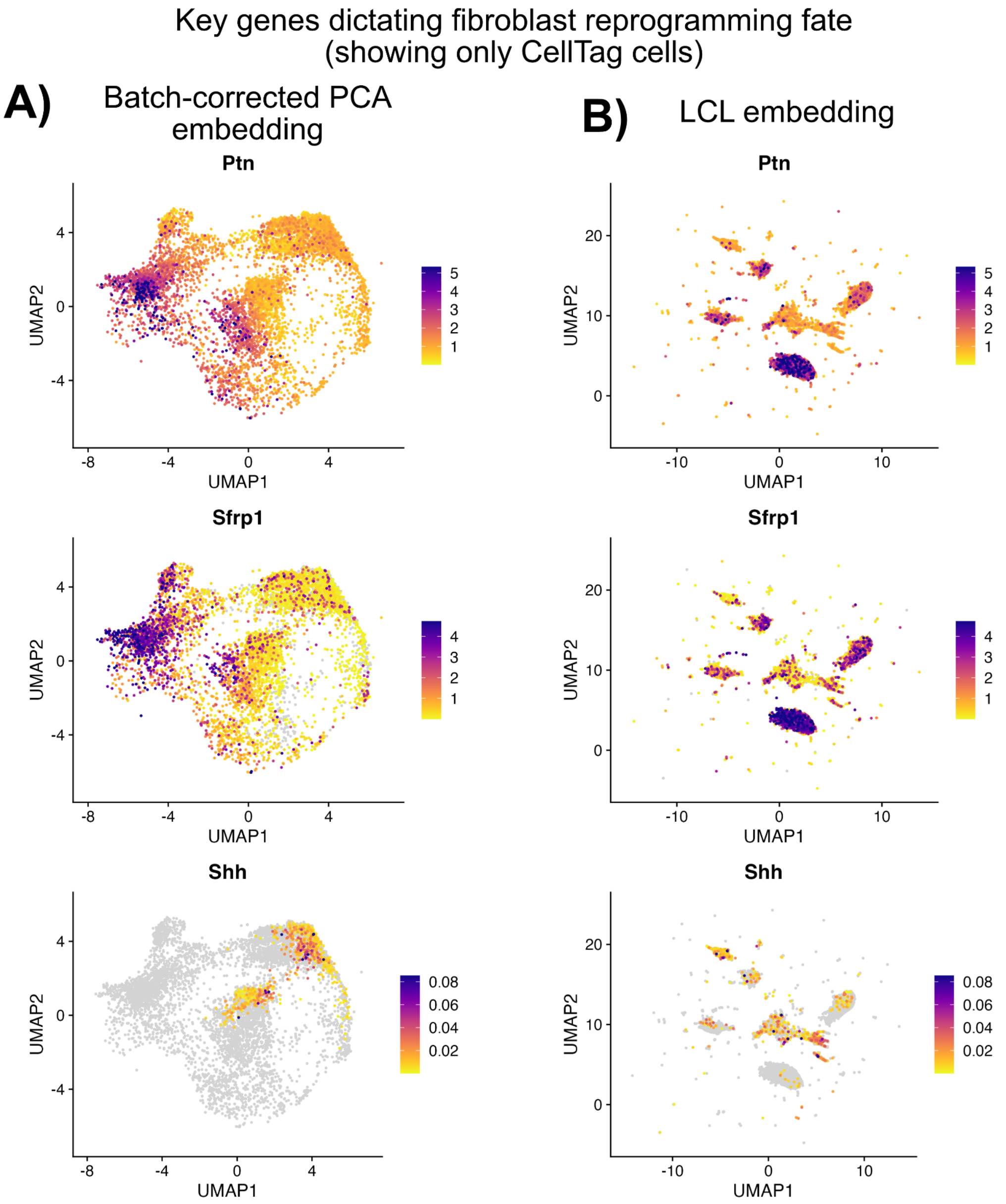


*Figure 17.* Key genes using the batch-corrected (original) embedding (**A**), compared to the LCL embedding (**B**). We observe more overt co-localization of key genes in the LCL embedding, illustrating its ability to cluster cells by lineage so that cells with similar fate dynamics share similar LCL coordinates.

*Table 11.* Dataset and Hyperparameter Configurations with Runtimes

| **Dataset** | **Num. cells** | **Num. cells per lineage** | **Hyper-parameters ($N$, $\alpha$, $U$, $\lambda$, $\tau$)** | **Runtime (hours)** |
|---|---|---|---|---|
| CellTag | 5893 | 34 | 50, 0.04, 5, 0.01, 0.5 | 1.1 |
| CellTag-multi | 22238 | 16 | 250, 0.02, 50, 0.01, 0.5 | 1.3 |
| CellTag(multi) Integration | 28131 | 19 | 250, 0.02, 50, 0.05, 0.5 | 1.3 |
| LARRY, Largest 200 | 10148 | 50 | 150, 0.5,15, 0.01, 0.5 | 1.02 |
| LARRY | 41093 | 14 | 260, 0.45, 15, 0.01, 0.5 | 1.78 |
| Pseudo-real, $\beta = 0.1$ | 41093 | 33 | 200, 0.5, 0, 0, 0.5 | 1.36 |
| Pseudo-real, $\beta = 0.3$ | 41093 | 42 | 150, 0.4, 0, 0, 0.5 | 1.09 |
| Pseudo-real, $\beta = 0.5$ | 41093 | 51 | 100, 0.4, 0, 0, 0.5 | 1.62 |
| Pseudo-real, $\beta = 0.7$ | 41093 | 68 | 100, 0.4, 0, 0, 0.5 | 2.04 |
| Pseudo-real, $\beta = 0.9$ | 41093 | 106 | 100, 0.35, 0, 0, 0.5 | 2.15 |
